# A Framework for Brain Atlases: Lessons from Seizure Dynamics

**DOI:** 10.1101/2021.06.11.448063

**Authors:** Andrew Y. Revell, Alexander B. Silva, T. Campbell Arnold, Joel M. Stein, Sandhitsu R. Das, Russell T. Shinohara, Dani S. Bassett, Brian Litt, Kathryn A. Davis

## Abstract

Brain maps, or atlases, are essential tools for studying brain function and organization. The abundance of available atlases used across the neuroscience literature, however, creates an implicit challenge that may alter the hypotheses and predictions we make about neurological function and pathophysiology. Here, we demonstrate how parcellation scale, shape, anatomical coverage, and other atlas features may impact our prediction of the brain’s function from its underlying structure. We show how network topology, structure-function correlation (SFC), and the *power* to test specific hypotheses about epilepsy pathophysiology may change as a result of atlas choice and atlas features. Through the lens of our disease system, we propose a general framework and algorithm for atlas selection. This framework aims to maximize the descriptive, explanatory, and predictive validity of an atlas. Broadly, our framework strives to provide empirical guidance to neuroscience research utilizing the various atlases published over the last century.

## Introduction

How we define anatomical brain structures and relate those structures to the brain’s function can either constrain or enhance our understanding of behavior and neurological diseases^1–4^. Discoveries by scientists like Carl Wernicke and Pierre Paul Broca, who mapped specific brain regions to speech function, in addition to case studies from Phineas Gage and H.M., who lost specific brain regions with resultant changes in brain function and behavior, exemplify how brain structure and function are fundamentally linked^5–7^. Properly labeling brain structures is paramount for enabling scientists to effectively communicate about the variability between healthy individuals and about the regions involved in neurological disorders^8^. Yet, no consensus has been reached on the most appropriate ways to label and delineate these regions, as evident by the wide variety of brain maps, or atlases, defining neuroanatomical structures^9^.

In common usage, an atlas refers to a “collection of maps”^10^ that typically defines geo-political boundaries and may include coarse borders (continental), fine borders (city), and anything in between (country; Fig. 1a, left). Borders^11^ are usually consistent across atlases of the world. In contrast, atlases of the brain are not consistent. Four separate atlases (Fig. 1a, right) may define the superior temporal gyrus differently. For example, approximately ninety percent of the *anterior* superior temporal gyrus in the Harvard-Oxford atlas^16^ overlaps with the *posterior* superior temporal gyrus in the Hammersmith atlas^17^. Atlases may also differ in other ways, including parcellation size, neuroanatomical coverage, and complexity of brain region shapes. For instance, the Yeo atlas^18^ contains 7 or 17 parcels while the Schaefer atlases^19^ may have between 100 and 1,000 parcels. Complicating matters further, atlases can differ in their intended use. The MMP atlas^20^ was intended for surface-based analyses^21^, yet a volumetric version (without subcortical structures) was independently created and used in connectivity studies^22^. The plethora of available atlases poses a problem for reproducibility in studying healthy and diseased populations and for metanalyses describing the involvement of different regions of the brain in various diseases. This has been termed the Atlas Concordance Problem^4^.

**Fig. 1.**
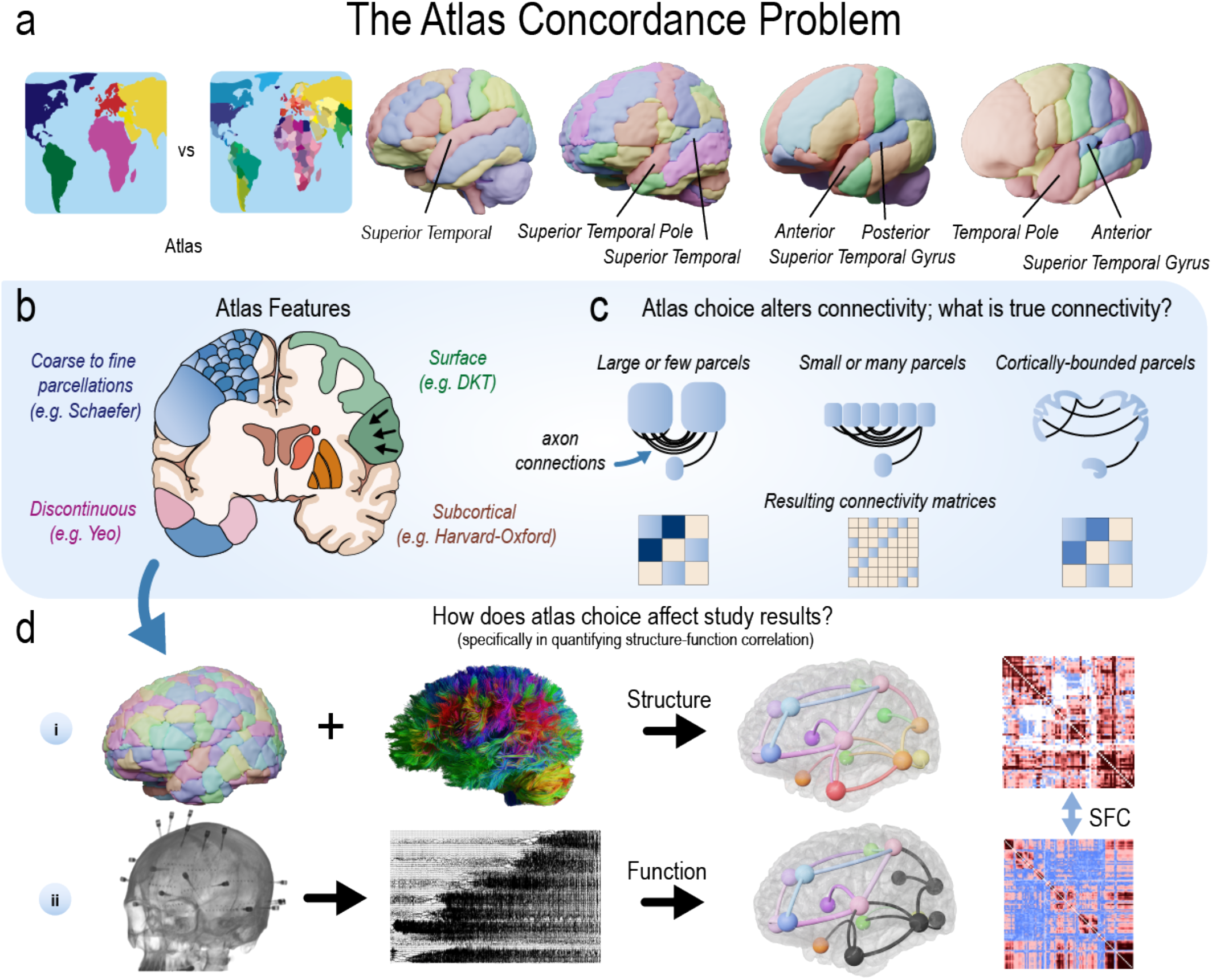
Many brain atlases are available in the neuroscience literature. **a**, In common usage, an atlas refers to a “collection of maps”^10^ that defines geo-political borders at different scales. Although borders^11^ are usually consistent across atlases of the world, they are typically not consistent across atlases of the brain. Four separate atlases (left-to-right: CerebrA, AAL, Hammersmith, Harvard-Oxford) may define the superior temporal gyrus differently. The lack of consistency across these labels poses a problem for reproducibility in cognitive, systems, developmental, and clinical studies, as well as metanalyses describing the involvement of different regions of the brain in various diseases^4^. This challenge has been previously referred to as the Atlas Concordance Problem. **b**, Atlases can have varying features (see also Table 1). **c**, Thus, all current connectivity studies in neuroscience may not accurately reflect some fundamentally “true” architecture. For example, atlases with either large or small parcels may affect the structural connectivity matrices that are used to define the “true” network architecture of the brain, and subsequently that are used to test hypotheses or make predictions about the brain. **d**, When combined with white matter tracts reconstructed from diffusion MRI, atlases can be used to measure how different regions of the brain are structurally connected (i). Similarly, intracranial EEG (iEEG) implants can record neural activity to measure how different regions of the brain are functionally connected (ii). Technologies such as fMRI, MEG, and many others can also measure functional connectivity. The statistical similarity between structural and functional connectivity measurements can be calculated (e.g., structure-function correlation; SFC). Such estimates have been used to better understand the pathophysiology of disease. In this study, we evaluate how the varying atlases may alter the power to test a specific hypothesis about the brain’s structure-function relationship in epilepsy.

In the present study, we perform an extensive evaluation of the available atlases in the neuroscience literature (Table 1) by examining the effect of varying features such as parcellation size, coverage, and shape (Fig. 1b) on structural connectivity (Fig. 1c). We also examine how atlas choice changes structural network topology by measuring structure-function correlation (SFC) using an atlas-independent measure of functional connectivity (Fig. 1d). We utilize a total of 55 brain atlases, including many routinely used in common neuroimaging software. Note the important distinction between the terms atlas, template, and stereotactic space^9^ (see Fig. S1). We found that different atlases may alter the *power* to test a hypothesis about epilepsy pathophysiology that seizures propagate through the underlying structural connections of the brain. This hypothesis has been previously supported in prior research^13,14,23,24^.

**Table 1.**
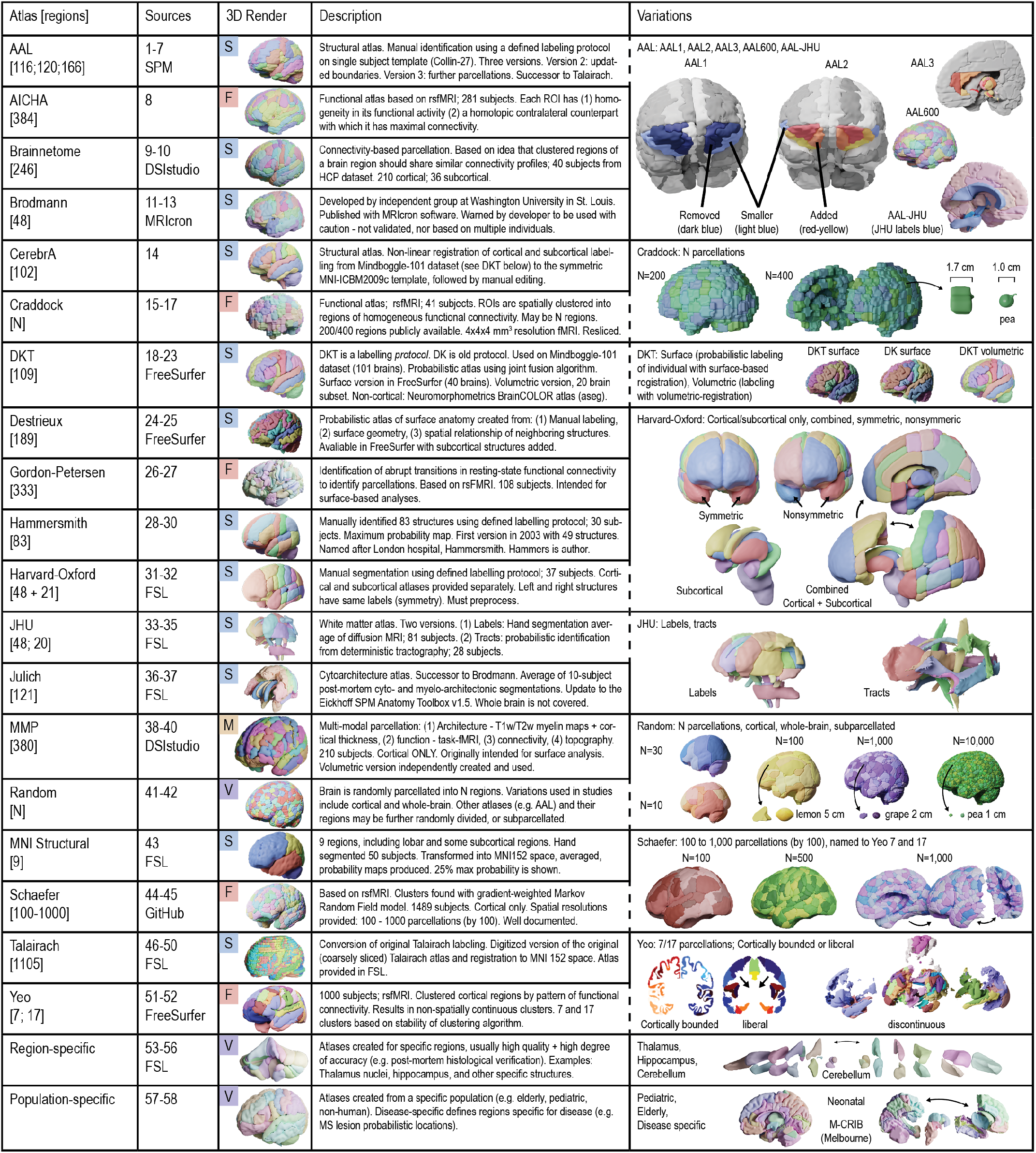
Atlases. Atlas sources are detailed in Table S1 and abbreviations are in the glossary. **S**: Structurally defined atlas; **F**: Functionally defined atlas; **M**: Multi-modally defined atlas; **V**: A variably defined atlas that may be structurally, functionally, or multi-modally defined; **ROI**: region of interest; **HCP**: Human connectome project dataset^12^; **MS**: multiple sclerosis.

In the context of our experimental design, we propose a new framework outlining how to appropriately choose an atlas when designing a neuroscience experiment. This framework is derived from historical foundations for assessing the validity and effectiveness of animal models^25^, network models^26^, and psychometric tests^27^, which try to maximize the (1) descriptive, (2) explanatory, and (3) predictive validity^26^ of a model. Atlases are a *tool* for investigators to test for causality and to make predictions about the brain. Thus, this framework incorporates a short discussion on explanatory modeling and predictive modeling, each with different goals (“To Explain or to Predict?”^15^). A one-size-fits-all approach may not exist for selecting an atlas, nor should it^28^; while there is one Planet Earth with a single atlas for a particular use (e.g., an atlas of the geo-political borders for a given point in time), there are many brains, with anatomical and functional variability across populations and species^28^. We hope our framework provides empirical guidance to neuroscience research utilizing the various atlases published over the last century.

## Results

### Clinical Data

Forty-one individuals (mean age 34 ± 11; 16 female) underwent High Angular Resolution Diffusion Imaging (HARDI), composed of thirteen controls (mean age 35 ± 13; 6 female) and twenty-eight drug-resistant epilepsy patients (mean age 34 ± 11; 12 female) evaluated for surgical treatment. Of the twenty-eight patients, twenty-four were implanted with stereoelectroencephalography (SEEG) and four with electrocorticography (ECoG). Ten SEEG patients (mean age 34 ± 8; 4 female) had clinical seizure annotations, and the first seizure from each patient (mean duration 81s) without artifacts was selected for SFC analyses. Patient and control demographics are included in Table S2.

### Atlas Morphology: Sizes and Shapes

We hypothesized that atlas morphological properties, including size and shape (Fig 2), affect SFC. To test this hypothesis, we first quantified the distributions of parcellation sizes (Fig 2a) and shapes (Fig 2b) in various atlases. These results exemplify the diversity of atlas parcellation morphology. Fig 2c shows a comparison of individual parcellation volumes and sphericities. The remaining atlases are shown in Fig. S2. In contrast to standard atlases, random atlases have constant sphericity with respect to volume size. Note that the distribution of parcellation shapes (i.e. sphericity) is similar across parcellation sizes in random atlases and their parcellations may not represent true anatomical or functional boundaries. Thus, random atlases allow us to study how parcellation scale affects network structure and SFC while keeping the effect of shape constant. Crucially, random atlases also allow us to explore if accurate and precise anatomical boundaries are essential in some experimental designs^29^.

**Fig. 2.**
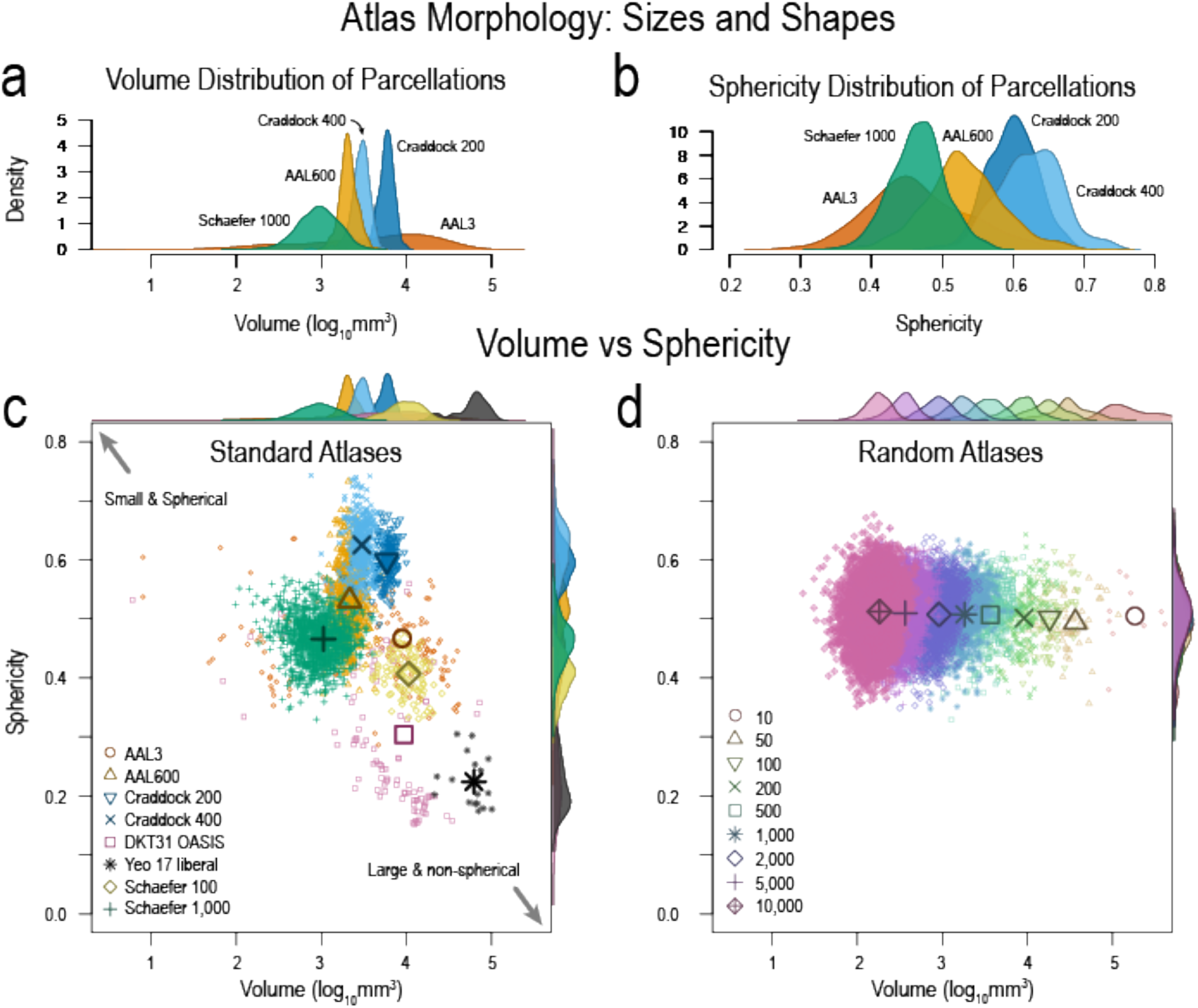
Atlas morphology: sizes and shapes. **a**, Volume distribution of atlas parcellations demonstrating the diversity of parcellation sizes. **b**, Parcellation sphericity distributions illustrating how the shapes of different parcellations may not be uniform. **c**, Volumes versus sphericity showing how some atlas parcellations may be small and spherical, while others may be large and non-spherical. This illustrates the non-uniformity in atlas parcellations. **d**, Volumes and sphericity of random atlases showing the uniformity of sphericity with changing volumes. Random atlases allow us to study (1) the effect of parcellation scale without the confound of shape effects and (2) the need for accurate anatomical boundaries to test a hypothesis about the structure-function relationship in the brain at seizure onset. Numbers in legend represent the number of parcellations for each random atlas. Remaining atlases are in Fig. S2.

### Varying atlases affect structural network topology

Although the morphology of atlas parcellations is diverse, we aimed to investigate how these morphological characteristics (particularly parcellation scale) affect structural network topology (Fig. 3). Networks are the basis upon which we compute SFC, and not necessarily morphological characteristics, therefore, we measured how network density, mean degree, characteristic path length, mean clustering coefficient, and small worldness change as a function of parcellation scale (Fig. 3a). We found that the change in these network measures are congruent between standard and random atlases and previous studies^30^. We also show that mean density, a global network measure, is similar between our control (N=13) and patient (N=28) cohorts (Fig. 3b).

**Fig. 3.**
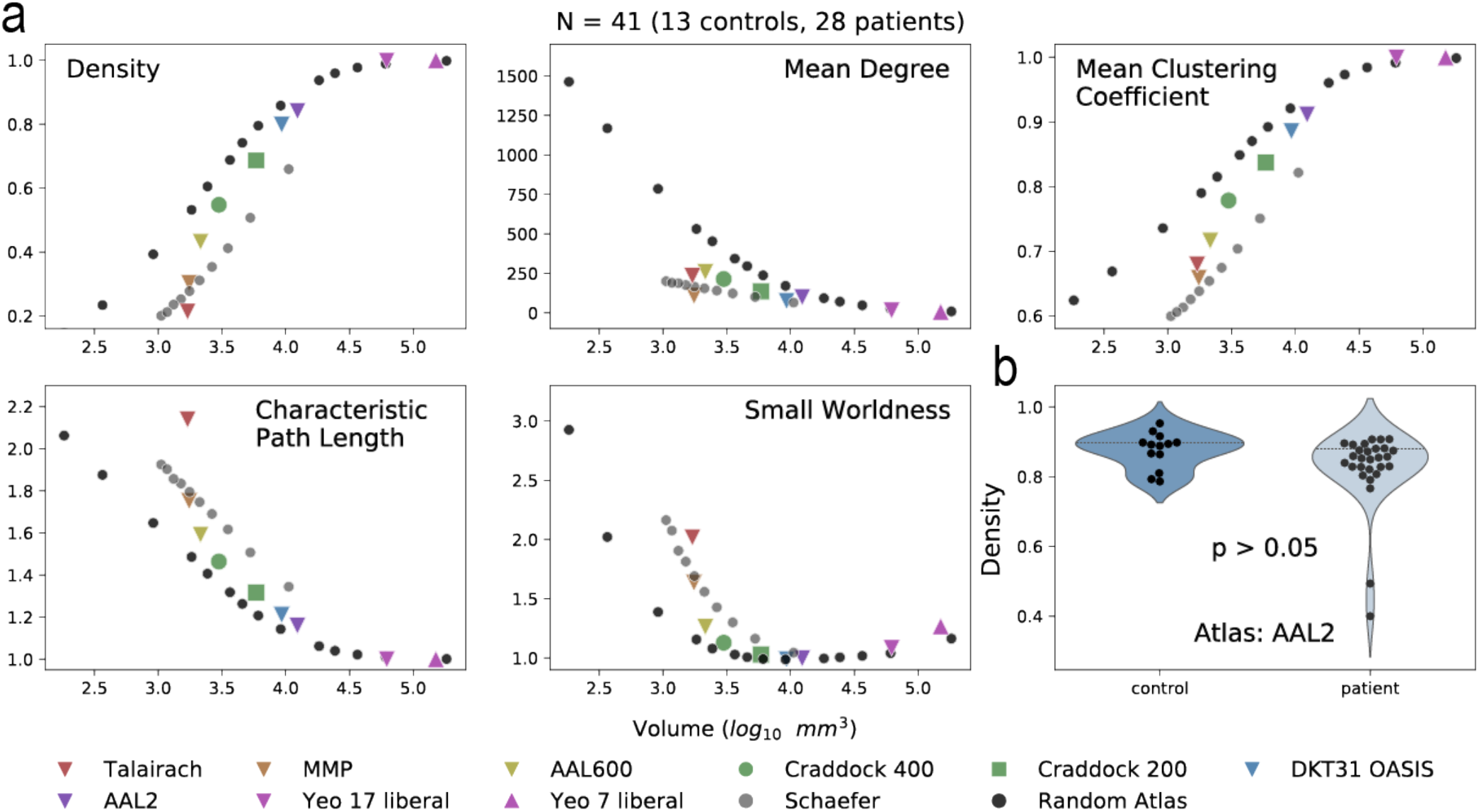
Structural network differences between atlases. **a**, Density, mean degree, mean clustering coefficient, characteristic path length, and small worldness were calculated for structural connectivity networks. A subset of atlases is shown. Remaining atlases studied are shown in Fig. S3. The average parcellation volume was calculated for each atlas and the corresponding network measure was graphed as the mean of all subjects (N=41; 13 controls, 28 patients). **b**, Controls and patients were not significantly different in density for the AAL2 atlas (Mann-Whitney U test), illustrating that global structural network measures are similar between cohorts. However, specific edge-level connections between cohorts may be different, and characterizing these differences is out of the scope of this manuscript. Controls and patients were separated and shown in Fig. S4. Network measures using different threshold are shown in Fig. S5.

### Varying atlases affect SFC: single subject

Fig. 4 illustrates an overview of how SFC is calculated. Structure is measured with high angular resolution diffusion imaging (HARDI) and function is measured with SEEG electrode contacts. Structural connectivity matrices are generated based on the atlas chosen (Fig. 4a) and functional connectivity matrices are generated based on broadband (1 – 127 Hz) cross-correlation of neural activity between the electrode contacts in widows of time (Fig. 4b, see Methods section on “Functional Connectivity Network Generation”). Thus, the structural network is static while the functional network is computed across time. The connectivity matrices shown are example data from a single patient, sub-patient07. Functional connectivity matrices are shown for 6 hours before seizure onset, 90 seconds before seizure onset (t = -90), 40 seconds after seizure onset (t = 40), 88 seconds after seizure onset (seizure duration = 89 seconds), and 180 seconds after seizure onset (91 seconds after seizure termination). Each functional connectivity matrix time window was correlated to each structural connectivity matrix, yielding a SFC at each time window (Fig. 4c). Each point represents the structural edge weight between two brain regions and their corresponding functional connectivity edge weight in broadband cross-correlation. A line of best fit is shown for visualization, and r values represent Spearman rank correlation for that time point. SFC was graphed for all time points during the interictal, preictal, ictal, and postictal periods for this patient in Fig. 4d.

**Fig. 4.**
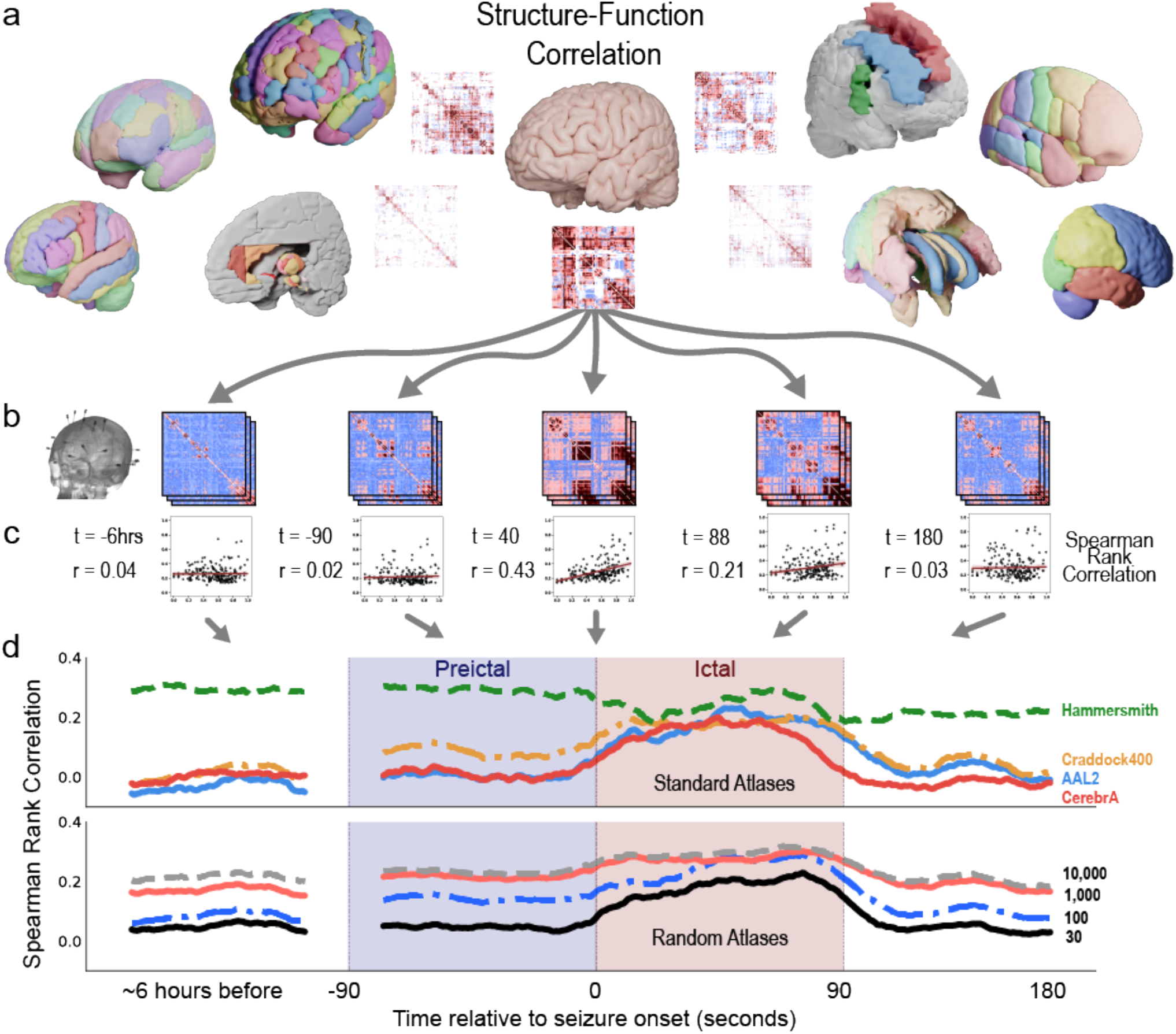
Structure-Function correlation in a single patient using different atlases. **a**, Example atlases and structural connectivity matrices. **b**, Functional connectivity matrices are computed from SEEG recordings during the interictal, preictal, ictal, and postictal periods. During each period, the SEEG data is binned into non-overlapping windows (the vertically stacked matrices) to create time varying representations of functional connectivity. Broadband cross correlation matrices are shown for sub-patient07 at 6 hours before seizure onset, 90 seconds before seizure onset, 40 seconds after seizure onset (t = 40), 88 seconds after seizure onset (seizure duration = 89 seconds), and 180 seconds after seizure onset (or 91 seconds after seizure termination). **c**, Each functional connectivity matrix is correlated to a structural connectivity matrix of a given atlas. Spearman Rank Correlation is measured between all time points and all atlases for each patient. Lines of best fit are for visualization purposes only. **d**, SFC is graphed at each time point for four example standard atlases (Hammersmith, Craddock400, AAL2, and CerebrA), and four example random atlases (30, 100, 1k, and 10k parcellations). SFC increases during seizure state for some standard atlases (Craddock 400, AAL2, and CerebrA atlases). This result follows previous SFC publications with ECoG^13,14^. However, SFC does not increase for the Hammersmith atlas. These findings highlight that the power to detect a change in the structure-function correlation at seizure onset, and thus the ability to probe the hypothesis that seizure activity is correlated to brain structure, may be reduced using some atlases. The use of different atlases may contradict previous studies.

Four example standard and random atlases are graphed. We show that SFC increases during the ictal state for many atlases (CerebrA, AAL2, Craddock 400), but not all atlases (Hammersmith). The increase in SFC during seizures follows previous SFC studies using ECoG^13,14^. Similarly, SFC increases for a subset of random whole-brain atlases. While parcellation scale may affect SFC, it is not the only feature affecting SFC – the Hammersmith and AAL2 atlases have similar parcellation scales yet diverging neuroanatomical properties and SFC dynamics. These findings highlight inference from one type of atlas may suggest that seizure activity is not correlated to brain structure, contradicting previous studies^13^.

### Varying atlases affect SFC: multiple subjects

Fig. 5 shows SFC for ten standard atlases and five random atlases using SEEG broadband cross-correlation metrics averaged across all patients with clinically annotated seizures (N = 10). The AAL2 atlas shows a statistically significant increase in SFC from preictal to ictal periods (p < 0.05 by Wilcoxon signed rank test after Bonferroni correction for 55 tests). This change from preictal to ictal SFC is denoted ΔSFC. Using the AAL2 atlas, this finding supports the hypothesis that seizure activity propagates and spreads via axon tracts making up the underlying structural connectivity of the brain^13,14^. SFC was similarly calculated for random whole-brain atlases. A notable finding is that during the interictal period, resting state SFC (rsSFC) increases at larger number of parcellations (i.e. smaller parcellation volumes). We show that rsSFC is observably affected by parcellation scale when plotting the random atlases in Fig. 5 (bottom row). These findings may be concerning given that the *inherent* structure-function relationship in the brain is not necessarily changing at resting state, but its measurement is greatly affected by atlas choice alone.

**Fig. 5.**
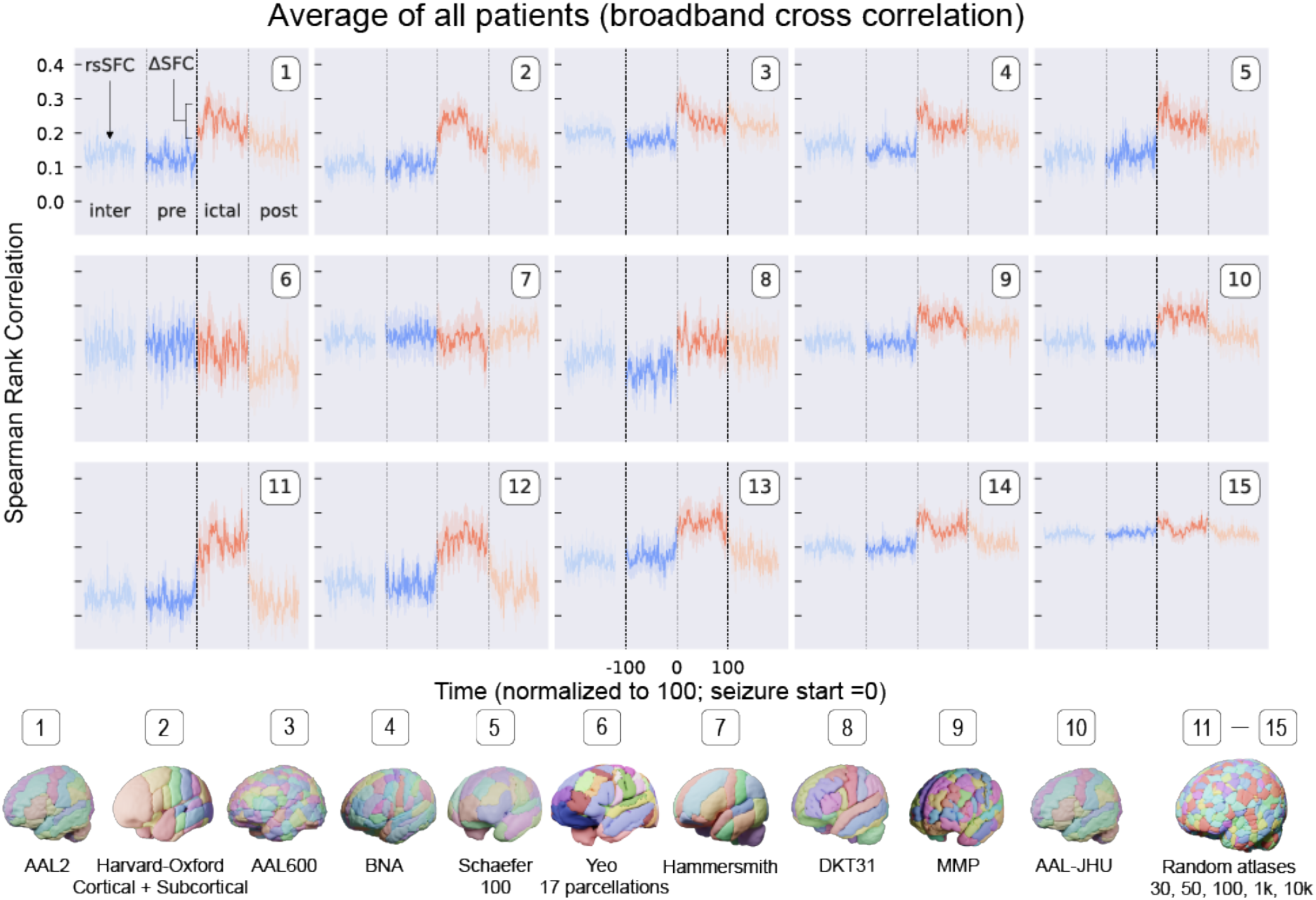
Structure-Function Correlation in multiple patients using different atlases. SFC for ten standard atlases and five random atlases using SEEG broadband cross-correlation matrices averaged across all patients with clinically annotated seizures (N = 10). Resting state SFC (rsSFC) is the SFC during the interictal period. The change from preictal to ictal SFC is ΔSFC. SFC was similarly calculated for random atlases and shows that rsSFC and ΔSFC may change with parcellation scale. These findings may be concerning given that the *inherent* structure-function relationship in the brain is not necessarily changing at resting state, but its measurement is greatly affected by atlas choice alone.

### Varying atlases affect resting state SFC and ΔSFC

Resting state SFC (rsSFC) and the change in SFC (ΔSFC) from preictal to ictal periods are affected by parcellation scale (Fig. 6). Fig. 6a shows how rsSFC *decreases* with larger average parcellation volumes (moving left to right). A large average parcellation volume for a given atlas generally means there is a fewer number of total parcellations (e.g. the MNI structural atlas has a large average parcellation volume given only nine parcellations). In contrast, Fig. 6b shows ΔSFC *increases* with larger parcellation volumes (moving left to right). Broadly, ΔSFC may be interpreted as the change in SFC with respect to a disease (e.g. a seizure) and non-disease states. This change metric has been used to characterize and make inferences in many neurological disorders^31,32^. Only a subset of atlases show a change in SFC at seizure onset (Fig. 6c). These results exemplify that either overly coarse or fine parcellations may not adequately capture the underlying SFC of the brain or its dynamics with relation to a neurological disease.

**Fig. 6.**
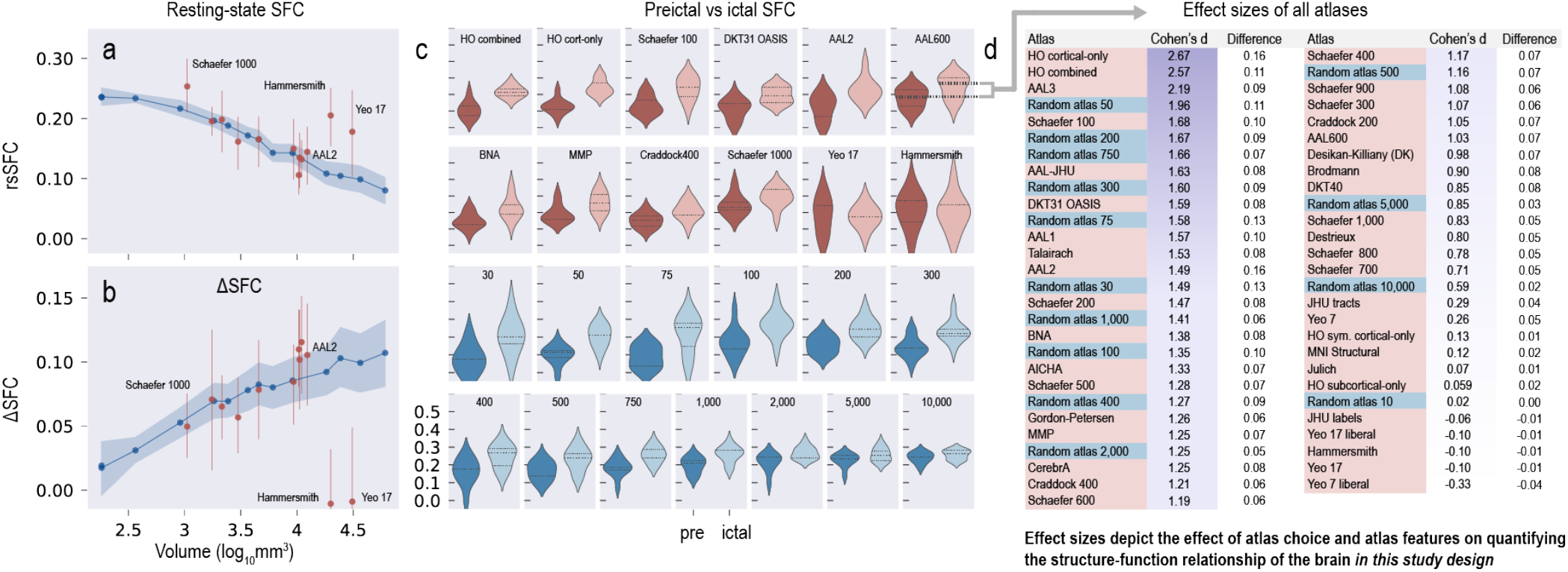
The power to test a hypothesis about epilepsy pathophysiology changes depending on atlas choice. **a**, Resting state SFC (rsSFC) decreases with larger parcellation volumes (moving left to right). Random atlases are shown in blue, and select standard atlases are shown in red. Points represent the average across all patients, and bands represent 95% confidence intervals. **b**, ΔSFC increases with larger parcellation volume (moving left to right). Broadly, |*Delta*SFC may be interpreted as the change in SFC with respect to disease (e.g. a seizure) and non-disease states, and this change has been used to characterize and make inferences on many neurological diseases. These results exemplify that parcellations that are either too coarse (large volumes) or too fine (small volumes) may not adequately capture the underlying SFC of the brain or its dynamics with relation to a neurological disease. **c**, A subset of atlases show a difference in preictal and ictal SFC. **d**, The effect size between preictal and ictal SFC is calculated for all 55 atlases used in this study. Many atlases commonly used in the neuroscience literature have comparable effect sizes to random atlases. The standard atlases with the greatest effect size (and thus power) are the Harvard-Oxford and AAL3 atlases. These atlases outperform many random atlases (where anatomical boundaries are not followed) and may indicate that their parcellation scheme captures the structure-function relationship in the brain at seizure onset with DTI and iEEG.

### Atlas choice affects the power to test a hypothesis

The effect size between preictal and ictal SFC is calculated for all 55 atlases used in this study (Fig. 6d). Cohen’s d and the difference between the mean ictal and mean preictal SFC are shown. Atlases are ordered by Cohen’s d.

We found that different atlases may alter the power to test the hypothesis about epilepsy pathophysiology that seizures propagate through the underlying structural tracts of the brain, measured with diffusion MRI. This hypothesis has been previously supported in prior studies^13,14,23,24^

Many atlases commonly used in the neuroscience literature have comparable effect sizes to random atlases (where anatomical boundaries are not followed). The standard atlases with the greatest effect size (and thus power, given equal significance levels and sample sizes) are the Harvard-Oxford and AAL3 atlases. These atlases outperform many random atlases and may indicate that their parcellations may adequately capture the structure-function relationship in the brain. These atlases may capture the “true” structural network architecture (see Fig. 1c) because these network architectures better differentiate and are more correlated to functional changes seen at seizure onset.

Despite the effect sizes of the Harvard-Oxford and AAL3 atlases, however, there may not be a “true gold standard” atlas or parcellation scheme given that resolution is more critical than the exact border location of parcels^29^, there may be no single functional atlas for an individual across all brain states^28^, and many standard atlases yield similar effect sizes to randomly generated atlases (this study).

## Discussion

In this study, we performed an extensive evaluation of the available structural, functional, random, and multi-modal atlases in the neuroscience literature (Table 1). We detailed morphological (Fig. 2) and network (Fig. 3) differences between these atlases. We showed the effect of atlas choice on the measurement of structure-function correlation (SFC) in epilepsy patients (Fig. 4 and Fig. 5). We also showed how various atlases may affect the power to test a hypothesis about seizure propagation (Fig. 6). This work has implications for investigators because the ability to test hypotheses and make predictions about the brain’s function may depend on atlas choice. In light of our study using an extensive list of available brain atlases, we propose a general framework below for evaluating and selecting an atlas (Fig. 7).

**Fig. 7.**
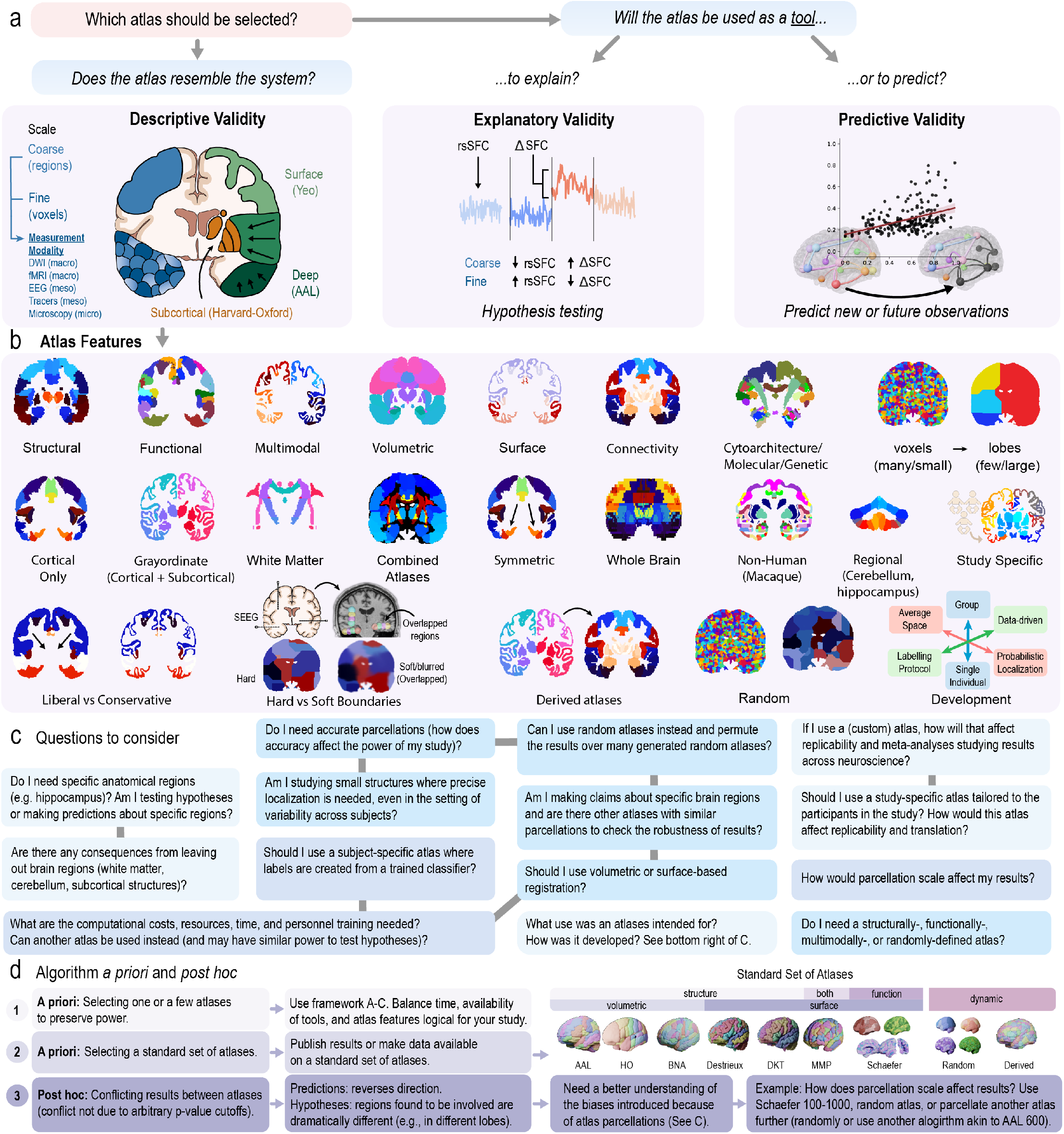
A Framework for brain atlases. **a**, Which atlas should be chosen for a study? We propose a framework that helps select an atlas in the context of its descriptive, explanatory, and predictive validity. **Descriptive validity** means the features of an atlas appropriately resembles the experimental system. An atlas is also a *tool* to solve a variety of problems in neuroscience. It may be used as part of a *methodology* to explain causality (**explanatory validity**), or it may be used to make predictions (**predictive validity**). These two goals are distinct, and the differences between explanation and prediction “must be understood for progressing scientific knowledge”^15^. These aspects (to explain or to predict) should be considered when selecting an atlas. **b**, Non-mutually exclusive atlas features related to descriptive validity. **c**, A list of questions to consider when choosing an atlas. Gray lines connect related questions. **d**, An algorithm for atlases selection *a priori* and *post hoc*. Please see the main text for further details.

### A Framework for Brain Atlases

Various publications have highlighted the Atlas Concordance Problem^2–4,9^, curated several atlases in freely accessible databases^33,34^, and made arguments for why specific atlas features (Fig. 7b) may be superior in certain situations^21,28,35–39^. There have been great efforts to publish accurate and precise parcellations as seen with an exponential rise in atlas-related publications over the last three decades (Fig. S8). However, none have found a general solution to the underlying problem: Does atlas choice matter?

We provide a framework that allows us to determine if the choice of an atlas is appropriate in the context of its (1) descriptive, (2) explanatory, and (3) predictive validity^26^. This framework is borrowed from the logic for assessing network models^26^, animal models,^25,40^, and psychometric tests^27,41^, where assessment of these models with standard statistical model-selection methods is particularly challenging. Thus, theoretical constructs already formulated in other fields may provide guidance.

### Descriptive validity

of an atlas refers to an atlas that appropriately resembles the system in which we work. In other words, it has “face value”^25^. An atlas should include features (Fig. 7b) relevant to the study (e.g., parcellations containing subcortical structures relevant to epilepsy). Importantly, the descriptive validity of an atlas also relates to the modality scale we use to measure the brain – for example, DWI and fMRI at the macroscale^42^, iEEG and tracers at the meso scale^43^, and microscopy at the microscale^44^. It is important to select a parcellation scale that resembles the measurement modality resolution (Fig. 6a). When correlating DWI with iEEG in our study at larger parcellation sizes, we lose our ability to discern precise anatomical locations that are structurally and functionally related (Fig. 6b). Similarly at smaller parcellation sizes (tending to voxel resolution), we may not capture the “true” structural network architecture (Fig. 1c), and thus we lose our ability to capture structure-function relationship changes at seizure onset.

An atlas is a *tool* to tackle a wide variety of problems in neuroscience. It may be part of a methodology to explain causality (**explanatory validity**) or it may be part of a methodology to make predictions (**predictive validity**). These two goals are distinct, and the differences between explanation and prediction “must be understood for progressing scientific knowledge” as described in “To Explain or Predict?” by Shmueli, 2010^15^. In the context of building scientific models, a model with a high explanatory ability may not have a high predictive ability.

Similar to models, atlases are also part of a scientific *methodology* to (1) explain how the brain functions or (2) predict new observations (i.e., they are one part of the overall methodological pipeline to test hypotheses or make predictions about the brain - for studies using atlases). Thus, atlases are tools. An atlas may be suitable for hypothesis testing, for example, because it includes subcortical structures like the hippocampus (also high descriptive validity) to support a hypothesis about seizure propagation through subcortical structures. Intuitively, without subcortical structures, it would be impossible to test hypotheses about subcortical structures. Less intuitively, explanatory validity of an atlas may also relate to the *power* to test hypotheses, which we show in our study. Some atlases may not be suitable for scientific inquiry because they provide little statistical power to detect differences in disease states, for example, to detect changes in SFC at seizure onset (Fig. 6b). It may be impossible to accurately predict power using an atlas before conducting a study, however, other studies asking similar questions using similar atlases may provide reasonable estimates of effect sizes (our study has similar effect sizes to a previous study^13^). Power may also depend on the accuracy of anatomical boundaries, or in our study, other atlas features such as parcellation scale and configuration (Fig. 6d). For example, the Harvard-Oxford and AAL3 atlases have similar parcellation configurations and similar power.

Some atlases may or may not be not be suitable for making predictions about new or future observations about the brain. For example, many network properties change with atlas choice (Fig. 3), and thus it is reasonable to suspect model prediction outputs may change with respect to the atlas used to build and train such models. Importantly, the exclusion of some anatomical structures, like white matter or the cerebellum in some atlases, may affect the training data used to build predictive models. In our study, a translational goal is to predict functional seizure activity from structural data. SEEG records activity from both gray matter and white matter; however, recent studies have shown that white matter functional recordings may provide different information than gray matter^45–48^. Thus, excluding some anatomical labels may affect model predictions. Another example is the use of network models to predict spread, such as *–*-synuclein across the brain connectome^49^. Without the incorporation of all brain structures related to *–*-synuclein spread, models to predict and monitor spread may be inaccurate.

### Are accurate anatomical or functional parcellations needed?

During the course of conducting this study, and while undergoing peer review, other atlases with more accurate or relevant parcellations to the study’s population were published in different areas of neuroscience^50–58^. Here, we cautiously propose a question: Are efforts to publish more atlases created with different algorithms or slightly modified parcellations from existing atlases providing any advantages over already existing atlases? Naturally, accurate and precise parcellations are needed when probing specific hypotheses about exact structures that depend on accurate segmentation of such structures (particularly at the sub-field or cellular level); however, few studies compare an atlas to a null atlas (one with randomly generated parcellations). Studies that do are Gordon et al. 2016^59^and Lewis et al. 2021^58^.

In this study, we show that random atlases provide similar power to detect differences in SFC between preictal and ictal states (Fig. 6d). Indeed, it is difficult or nearly impossible to evaluate a newly proposed atlas, given that the performance metrics to evaluate an atlas may be infinite (given infinite experimental designs). Only one such metric, SFC, was used in this study. But given new deep learning methods and other computationally expensive methods using trained classifiers for segmentation, existing atlases may be adequate for labs with limited funding resources, trained personnel, and access to GPUs. These labs may still be capable of answering important questions in neuroscience.

### Which atlas should be used for my study?

One of the most difficult challenges as scientific investigators is to make optimal methodological decisions to discover useful findings for the scientific community. Selecting an atlas is one such decision we may make in some of our studies. We realize the framework provided above may be abstract to some readers; we also provide a concrete list of questions to consider when choosing an atlas (Fig. 7c) for a neuroimaging study. However, in conducting this study, we also found that researchers may face three problems when choosing an atlas (Fig. 7d) and these problems are worth further discussion. The first two problems are in selecting an atlas *a priori*, or before conducting a study. They deal with selecting one or a few atlases to preserve power, or in selecting a standard set of atlas to publish public data for other researchers to use. The third problem is the issue of conflicting results between two atlases and what to do after a study is conducted (*post hoc*). We provide a further discussion on these problems below.

### Considerations in selecting one or a few atlases

Selecting one atlas may preserve power and avoid a multiple comparisons problem by testing every atlas. Selecting an additional atlas may also be chosen to confirm the robustness of results. In these cases, a balance of time, availability of tools, and atlas features logical for your study as outlined in Fig. 7a-c need to be considered. For example, if a custom atlas is used, how will that affect replicability and meta analysis in the long-run for the field? What are the atlas features needed (such as scale and coverage of regions)? What are the computational costs and personnel training needed to use particular atlases? (See questions in Fig. 7c).

### Considerations in selecting a standard set of atlases

When publishing results and/or making data publicly available for other investigators to use, another approach is to select a set of atlases based on the perceived needs of other investigators, atlas features covered, prevalence of atlases used in the literature (Fig. S9a), and the prevalence of “turn-key” neuroimaging software that incorporate these atlases (Fig. S9b). Studies are emerging with data publicly available for use based on one or a few select atlases^60,61^. Many turn-key neuroimaging software also inevitably have to make the decision to employ a set of atlases to meet the needs of many researchers. A problem may arise, however, when other researchers need the published data at other atlas resolutions or with other structures. And unfortunately, the value of the data may be lessened and the effort put in by the publishing researchers may be in waste if this happens. What may help with the atlas concordance problem is perhaps a “standard set” of atlases – a set to benchmark studies across the neuroimaging field. Furthermore, turn-key tools like FreeSurfer, QSIprep, DSI-studio, FSL, and many others may benefit from a standard set of incorporated atlases that captures enough features useful to the majority of the neuroscience community, even if not every available atlas is included. Based on our exhaustive search of atlases in the neuroimaging literature, the ability to collect them for use in a single study, the prevalence of certain atlases already in-use (Fig. S9a), and the prevalence of neuroimaging software (Fig. S9b) we propose an initial set of atlases (Fig. 7d).

The AAL atlas is one of the most commonly used volumetric atlases (Fig. S9a), and along with the Harvard-Oxford atlas, may provide complimentary results when published together. The Brainnetome atlas^62^ is another structural atlas at a finer resolution, having gained popularity since its introduction in 2016. The Destrieux and DKT atlases are also structural atlases, and already incorporated into one of the most commonly used neuroimaging software, FreeSurfer (https://surfer.nmr.mgh.harvard.edu). FreeSurfer provides surface-based registration, which may more accurately label cortical structures than volumetric registration (Fig. S6). Accurate segmentation of sub-cortical structures may also be acquired from FSL^63^ (https://fsl.fmrib.ox.ac.uk/fsl/fslwiki). In addition, the MMP, or “Glasser” atlas was created from multi-modal imaging data. A commonly used atlas provided at different scales is Schaefer atlases provide, however, it does not include subcortical structures.

Random atlases may also provide robust conclusions by allowing researchers to manipulate the resolution, size, and shape of parcellations and iterate over many atlases. Although random parcellations may forgo accuracy because they do not follow true anatomical boundaries, these atlases may still provide similar conclusions to other standard atlases with the added benefit of permuting results over many atlases (Fig. 6). An alternative to random atlases is to divide or combine the parcellations of another standard atlas (a “derived” atlas in Fig. 7d. For example, the AAL 600 is derived from the AAL atlas in which its parcellations are further sub-divided using a specified algorithm. Parcellations may also be sub-divided randomly.

### Considerations in conflicting results between atlases

When more than one atlas is used, results may conflict. We define conflicting results as two different atlases giving alternating predictions (e.g., good vs poor outcomes, increase in SFC rather than decrease in SFC) or support alternating working hypotheses (e.g., the temporal lobe is involved in one atlas, but another atlas highlights the involvement of the frontal lobe in the pathophysiology of a disease). We do not mean that conflicting results arise due to lack of statistical power (e.g., one atlas gives a p-value of 0.06 and another atlas 0.04).

One way to understand if the observed effect is not an artifact of the atlas choice is to select a few atlases with varying features and figure out what is causing the conflict. Unfortunately, there may be no other way given that every study will have different parameters and measurements to know what gives rise to conflicting results. In the matter where conflicting results arise due to atlas selection, then it may troubleshooting may be needed to understand what gives rise to the conflict (surface vs volumetric registration, parcellation scale, missing relevant structures, etc.). Fortunately, however, most atlases in this study affect power rather than conflicting results (Fig. 6d. We hope this discussion, our study, and our figures provide insight to others.

## Limitations

Our study is not without limitations. A major limitation is that we did not evaluate atlases in a diverse set of experimental systems, but rather limited our analysis to a contemporary topic in epilepsy using SEEG implantations and to a study of the structure-function of the brain, potentially appealing to a wider audience. The question we were trying to answer (“Which atlas should we use?”) is a difficult problem to solve, given that it would be impossible to evaluate all atlases in all experimental designs. We attempted to generalize a framework given our findings after an extensive search for, and curation of, available neuroimaging atlases.

We also did not perform a feature selection analysis posthoc to maximize ΔSFC at seizure onset; rather, we performed a comprehensive evaluation of many atlases to set a general framework and describe the nuances between the different atlases and their features. Ideally in our study, we required a whole-brain, volumetric atlas that covered the implanted SEEG electrode contacts. No such atlas existed. We opted for combining different atlases or developing randomly parcellated atlases used in previous publications^30,64^. However, no general framework existed to determine which atlas should be used or clearly outlined the feature space of these atlases. We had no formal basis for how changing an atlas could change our results and eventual goal for translating network models to better treat epilepsy patients.

Another limitation, we assume a change in SFC supports the hypothesis that seizures harness the underlying structural connectome of the brain (along with support from prior literature^13,14,65^). We may be biasing our results to select an atlas that maximizes ΔSFC. However, we wish to select a methodology that allows us to measure *any change* in brain state that accompanies seizure onset (explanatory validity), permitting us to probe epilepsy biology and understand the processes that govern seizure spread.

An additional limitation concerns the effect of parcellation volume on SFC. In probing this effect across our random atlases and atlases used in the literature, we did not perform controlled experiments to separate the effects of parcellation size from parcellation N (number of parcellations). A future experiment could fix the number of parcellations while changing parcellation volume (or vice versa). This would allow us to test whether parcellation volume or N drives changes in SFC. However, this was outside the scope of our study.

Our goal was to highlight the importance of selecting an appropriate atlas from an array of possibilities, using a datadriven, validated experimental paradigm^13^. We acknowledge new studies that show that streamline counts may not completely reflect the underlying diffusion data^66^; however, comparing such techniques were outside the scope and goal of our focused study. We also note that few patients had lesions on imaging. Misalignment due to non-linear distortion may add noise to our data; however, few patients had lesions. Our study was not conducted to necessarily make the claim that SFC changes exist in the brain at seizure onset, but rather to show how varying atlases may change SFC.

Finally, our analysis relies on the assumption that an atlas approach must be used to quantify SFC and does not consider an atlas-agnostic approach nor if such an approach is appropriate. To study SFC using networks, both structural and functional networks must have nodes representing the same entity – neuroanatomical structures. The atlases defining anatomical structures (whether they are functionally, histologically, genetically, procedurally, multi-modally, or randomly defined) are the link between structural connectivity and functional connectivity measurements of the brain. To study SFC, we must rely on the neuroanatomical structures defined by an atlas, then localize electrodes to these regions and correlate the structural measurements (e.g., streamlines, fractional anisotropy, mean diffusivity) with functional measurements (e.g., cross-correlation, coherence, mutual information). Fundamentally, we are defining the nodes of the brain in advance, which can alter our results; a more comprehensive discussion on defining the nodes of the brain are in Fornito et al., 2016 and Bijsterbosh et al., 2017^43,67^.

## Conclusion

The publication of atlases and their distribution across neuroimaging software platforms has risen exponentially over the last three decades. Our study illustrates the critical need to evaluate the reproducibility of neuroscience research using atlases published alongside tools and analysis pipelines already established in the neuroscience community (e.g., FreeSurfer, DSI studio, FSL, SPM, QSIprep, fMRIprep, MRIcron, ANTs, and others).

## Materials and Methods

### Human Dataset

MRI data was collected from forty-one individuals, including thirteen healthy controls and twenty-eight drug-resistant epilepsy patients at the Hospital of the University of Pennsylvania. Twenty-four patients underwent stereoelectroencephalography (SEEG) implantation and four underwent electrocorticography (ECoG) implantation. Ten of the SEEG patients had clinically annotated seizures and were used for SFC analyses. Inclusion criteria consisted of all individuals who agreed to participate in our research scanning protocol, and (if they had implantations) allowed their de-identified intracranial EEG (iEEG) data to be publicly available for research purposes on the International Epilepsy Electrophysiology Portal (https://www.ieeg.org)^68,69^. Seizure evaluation was determined via comprehensive clinical assessment, which included multimodal imaging, scalp and intracranial video-EEG monitoring, and neuropsychological testing. This study was approved by the Institutional Review Board of the University of Pennsylvania, and all subjects provided written informed consent prior to participating. See Table S2 for subject demographics.

### Structure

Methods and pipelines for structural connectivity generation and analysis are described in the following sections. Specific GitHub files and code are included where applicable.

### Imaging Protocol

Prior to electrode implantation, MRI data were collected on a 3T Siemens Magnetom Trio scanner using a 32-channel phased-array head coil. High-resolution anatomical images were acquired using a magnetization prepared rapid gradient echo (MPRAGE) T1-weighted sequence (repetition time = 1810 ms, echo time = 3.51m, flip angle = 9, field of view = 240mm, resolution = 0.94×0.94×1.0 mm3). High Angular Resolution Diffusion Imaging (HARDI) was acquired with a single-shot EPI multi-shell diffusion-weighted imaging (DWI) sequence (116 diffusion sampling directions, b-values of 0, 300, 700, and 2000s/mm2, resolution = 2.5×2.5×2.5 mm3, field of view = 240mm). Following electrode implantation, spiral CT images (Siemens) were obtained clinically for the purposes of electrode localization. Both bone and tissue windows were obtained (120kV, 300mA, axial slice thickness = 1.0mm)

### Diffusion Weighted Imaging (DWI) Preprocessing

HARDI images were subject to the preprocessing pipeline, QSIPrep, to ensure reproducibility and implementation of the best practices for processing of diffusion images^70^. Briefly, QSIPrep performs advanced reconstruction and tractography methods in curated workflows using tools from leading software packages, including FSL, ANTs, and DSI Studio with input data specified in the Brain Imaging Data Structure (BIDS) layout.

### Structural Network Generation

DSI-Studio (http://dsi-studio.labsolver.org, version: December 2020) was used to reconstruct the orientation density functions within each voxel using generalized q-sample imaging with a diffusion sampling length ratio of 1.25^71^. Deterministic whole-brain fiber tracking was performed using an angular threshold of 35 degrees, step size of 1mm, and quantitative anisotropy threshold based on Otsu’s threshold^72^. Tracks with length shorter than 10mm or longer than 800mm were discarded, and a total of 1,000,000 tracts were generated per brain. Deterministic tractography was chosen based upon prior work indicating that deterministic tractography generates fewer false positive connections than probabilistic approaches, and that network-based estimations are substantially less accurate when false positives are introduced into the network compared with false negatives^30^. To calculate structural connectivity, atlases listed in Table 1 were used. Structural networks were generated by computing the number of streamlines passing through each pair of structural regions in each specific atlas. Streamline counts were log-transformed and normalized to the maximum streamline count, as is common in prior studies^24,73−75^. GitHub: packages/imaging/tractography/tractography.py

### Atlases

Atlas descriptions and sources used in this study are found in Table S1. The 55 atlases used are listed explicitly in the reporting of effect sizes in Fig. 7d. All atlases were sourced in MNI space and if not already, resliced to dimensions 182×218×182. Atlases were linear and non-linear registered to T1w subject space using the ICBM 2009c Nonlinear Asymmetric template^76^ and FSL flirt and fnirt^77^.

We also included three atlases registered using surface-based approaches. These atlases (the DKT, DK, and Destrieux atlases) are output from FreeSurfer’s recon-all pipeline^78^. Many neuroimaging studies and software use volumetric approaches for registration^21^, yet surface-based approaches may yield more accurate labeling of the cortical surface (Fig. S6). The DKT40 atlas referred in this study is the surface version, while the DKT31 OASIS is the publicly available volumetric version (see Table S1).

In addition to published standard atlases above, we used whole-brain random atlases. A limitation of standard atlases is that they may not have anatomical definitions for all regions of the brain, and therefore, implanted electrodes may not be assigned properly to a region. This limitation was the impetus of our study (i.e., selecting an appropriate atlas for SEEG electrode localization and quantifying SFC). Whole-brain random atlases, in contrast, provide coverage to all implanted electrodes. They allow for the ability to change some morphological properties (i.e. parcellation size), while keeping other morphologies the same (e.g., parcellation shape; Fig. 2d). However, a limitation of random atlases is that their regions may not represent true anatomical or functional boundaries. Random atlases were built in the ICBM 2009c Nonlinear Asymmetric template space and covered all voxels, excluding those labeled as CSF or outside the brain. To fill these points, a pseudo grassfire algorithm was applied^30^. Briefly, N points representing the number of parcels of the atlas were randomly chosen as seed points. These seed points were iteratively expanded in all six Cartesian directions until all points were covered by one of the initial N seeds. After each iterative step, the smallest volume region expanded first. Random atlases created were of N equal to 10, 30, 50, 75, 100, 200, 300, 400, 500, 750, 1000, 2000, 5000, and 10000 parcels. Five permutations for each N were created. GitHub code to generate random atlases: packages/imaging/randomAtlas/randomAtlasGeneration.py

### Atlas Morphology: Volume and Sphericity

Atlas morphological measurements included parcellation size (volume) and shape (sphericity) (Fig. 2). Parcellation volume was calculated as the number of voxels in an parcel and log10 transformed. Parcellation sphericity was calculated as the ratio of the surface area of a sphere with an equal volume of the parcellation to the actual surface area of the atlas parcellation. Under this definition, sphericity is bounded from 0 to 1 where 1 is a perfect sphere. For reference, a perfect cube and a hemi-sphere have a sphericity of 0.8 and 0.7 respectively. GitHub: packages/imaging/regionMorphology/regionMorphology.py

### Structural Network Measures

We characterized the structural network topology of 52 atlases (Fig. 3 and Fig. S3). The three surface-based atlases (DKT40, DK, and Destrieux atlases output from the FreeSurfer recon-all pipeline^78^) were excluded from analyses of Fig. 2 and Fig. 3 because they were individually registered to each subjects’ T1w image. To quantify network topology, we examined density, mean degree, mean clustering coefficient, characteristic path length, and small worldness. Connectivity matrices were first binarized, using a threshold of 0, and a distance matrix was computed. The same binarization process and threshold was used across all atlases. The distance of any nodes that were disconnected from the main graph was set to the maximum distance between any pair of nodes in the main graph. Density, mean degree, clustering coefficient, and characteristic path length were then calculated on the binary, undirected graphs. Small worldness was calculated as the σ-ratio where σ = γ/λ and is the ratio of the average, normalized clustering coefficient, C, to the normalized characteristic path length, I. γ = CG/CR and λ = lG/lR where G is the graph of interest and R represents a ‘random’ graph that is equivalent to G. To approximate the equivalent random graph R due to intractable computational costs^79^, a well-known analytical equivalent CR = d/N and IR = log N/log d were used, where d denotes average nodal degree. All network measures were calculated using the Brain Connectivity Toolbox for Python. GitHub: papers/brainAtlas/Script_05_structure_02_network_measures.py

### Function

Methods and pipelines for functional connectivity generation and analysis are described in the following sections. Specific GitHub files and code are included where applicable.

### Intracranial EEG Acquisition

Stereotactic Depth Electrodes were implanted in patients based on clinical necessity. Continuous SEEG signals were obtained for the duration of each patient’s stay in the epilepsy monitoring unit. Intracranial data was recorded at either 512 or 1024 Hz for each patient. Seizure onset times were defined by the unequivocal onset^80^. All annotations were verified and consistent with detailed clinical documentation. If a patient had more than one seizure annotated, the first seizure longer than 30 seconds without artifacts was used.

### Electrode Localization

In-house software^81^ was used to assist in localizing electrodes after registration of pre-implant and post-implant neuroimaging data. All electrode coordinates and labels were saved and matched with the electrode names on IEEG.org. All electrode localizations were verified by a board-certified neuroradiologist (J.S.). Electrode contact assignment to atlas region assignment was performed by rounding electrode coordinates (x,y,z) to the nearest voxel and indexing the given atlas at that voxel in the same space as the patient’s T1w image. Electrodes that fell outside the atlas of interest were excluded from subsequent analysis. Please see Fig. S10 for visualization. We also show the percentage of contacts assigned a region given an atlas (Fig. S7) GitHub: packages/atlasLocalization/atlasLocalization.py

### Functional Connectivity Network Generation

Functional connectivity networks were generated from four periods: interictal, preictal, ictal, and postictal. (1) The interictal period consisted of the time approximately 6 hours before the ictal period. (2) The preictal period consisted of the time immediately before the ictal period. (3) The ictal period consisted of the time between the seizure unequivocal onset and seizure termination. (4) The postictal period consisted of the time immediately after the ictal period. Interictal, preictal, and postictal periods were 180 seconds in duration. Following removal of artifact-ridden electrodes, SEEG signals inside either GM or WM for each period were common-average referenced to reduce potential sources of correlated noise^82^. Next, each period was divided into 2s time windows with 1s overlap^83–86^. To generate a functional network representing broadband functional interactions between SEEG signals (Fig. 4b), we carried out a method described in detail previously^13,85^. Namely, signals were notch-filtered at 60 Hz to remove power line noise, low-pass and high-pass filtered at 127 Hz and 1Hz to account for noise and drift, and pre-whitened using a first-order autoregressive model to account for slow dynamics. Functional networks were then generated by applying a normalized cross correlation function *ρ* between the signals of each pair of electrodes within each time window, using the formula:

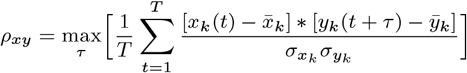

where x and y are signals from two electrodes, k is the 2s time window, t is one of the T samples during the time window, and *τ* is the time lag between signals, with a maximum lag of 0.5 s. Here, *σ* represents the standard deviation of the signal. Note that functional connectivity measurements were also calculated for coherence and zero time-lag Pearson and Spearman rank correlations with associated p-values in defined frequency bands reviewed in Newson and Thiagarajan 2019^87^, but were not analyzed or used in hypothesis testing in the study. For data, available data, please see “Data availability and Reproducibility” section below. Networks are represented as fully-weighted connectivity matrices. GitHub Code: GitHub: code/tools/echobase.py

### Structure-Function Correlation

To quantify the relationship between structure and function in the epileptic brain, we computed the Spearman rank correlation coefficient between the edges of the structural connectivity network and the edges of the functional connectivity networks (Fig. 4c). To avoid redundancy given the symmetric nature of the matrices, only the upper triangle was analyzed. In brief, the structural connectivity network, representing normalized streamline counts between each atlas region, was first down sampled to only include regions that contained at least one SEEG contact Fig. S10. This gave one static representation of structural connectivity. In the case where multiple electrodes fell in the same atlas region, a random electrode was selected to represent the functional activity of that neuroanatomically defined region. Next, for every time-window of the functional network, the functional network edges were correlated with the down sampled, static structural network edges. This resulted in a structure-function correlation time series. Note that atlases with very small region volumes included more electrodes for SFC calculation. Electrodes that did not localize to an atlas were excluded from analysis. To average the SFC for all patients and each atlas (Fig. 5), SFC time-series was resampled to 100 seconds for each period and each sample was averaged together. GitHub code: packages/eeg/echobase/echobase.py

### rsSFC and ΔSFC

Resting-state SFC (rsSFC) was defined as the SFC during the interictal period, approximately 6 hours before the ictal period. The mean SFC of that period was computed. ΔSFC was defined as the change in the mean SFC from the preictal to the ictal period (Fig. 5 top left panel). rsSFC and ΔSFC was calculated for each atlas (Fig. 6).

### Statistics

Preictal and ictal SFC for each atlas were compared using effect sizes across the 55 atlases shown in Fig. 6d. Cohen’s d and the difference between preictal and ictal SFC was calculated.

### Data availability and Reproducibility

All code files used in this manuscript are available at https://github.com/andyrevell/revellLab. All de-identified raw and processed data (except for patient MRI imaging) are available for download by following the links on the GitHub. Raw imaging data is available upon reasonable request from Principal Investigator K.A.D. iEEG snippets used specifically in this manuscript are also available, while full iEEG recordings are publicly available at https://www.ieeg.org. The Python environment for the exact packages and versions used in this study in contained in the environment directory within the GitHub. The QSIPrep docker container was used for DWI preprocessing.

## Acknowledgements

We thank Adam Gibson, Carolyn Wilkinson, Jacqueline Boccanfuso, Magda Wernovsky, Ryan Archer, Kelly Oechsel, members of Andrew’s Thesis Committee, and all other members and staff of the Center for Neuroengineering and Therapeutics for their continued help and support in this work.

## Supplementary Material

Please see supplemental figures and tables contained below.

- Figures
  Fig. S1: Atlas, Template, and Coordinate (Stereotactic) Space
  Fig. S2: Atlas Morphology: Sizes and Shapes (All atlases)
  Fig. S3: Network measures for remaining atlases
  Fig. S4: Network measures for controls and patients separated
  Fig. S5: Network measures for different thresholds
  Fig. S6: Effects of Registration: Volumetric- and Surface-based approaches
  Fig. S7: Coverage of electrode contacts
  Fig. S8: “Brain Atlas” Search in PubMed
  Fig. S9: Prevalence of select brain atlases and neuroimaging software
  Fig. S10: Electrode localization and region selection
- Tables
  Table. S1: Atlas Sources and References (3 pages).
  Table. S2: Patient and Control Demographics
- Other materials
  Glossary

## Glossary

1. **Atlas abbreviations and definitions**. For further details, see Table. S1.
  a. **AAL**. Automated anatomical labeling atlas.
  b. **AAL1, AAL2, AAL3**. AAL atlas versions 1, 2, and 3, respectively.
  c. **AAL-JHU**. The AAL atlas and the JHU labels atlas combined. For overlapping regions, the JHU atlas takes precedence.
  d. **AAL600**. AAL atlas with 600 parcels.
  e. **AICHA**. Atlas of Intrinsic Connectivity of Homotopic Areas.
  f. **BNA**. Brainnetome atlas.
  g. **Craddock 200-400**. Craddock atlases with a specified number of parcels (e.g. Craddock 200 will have 200 parcels). There are two atlas sizes publicly available - the Craddock 200 and Craddock 400 atlases.
  h. **DKT31 OASIS**. The DKT atlas from the OASIS dataset. See Table. S1 sources for more details. It is the volumetric version.
  i. **DKT40**. The DKT atlas used as part of FreeSurfer. See Table. S1 sources for more details. It is the surface version.
  j. **DK**. The Desikan-Killiany atlas. Surface atlas from FreeSurfer.
  k. **HO**. Harvard-Oxford atlas.
  l. **HO cortical-only**. HO atlas with only cortical regions. The symmetrical regions (the same region name on the contralateral hemisphere) are labeled with *different* identifications. Thus, this atlas has *non-symmetrical* labels (e.g. both temporal pole regions are labeled with a different identification number). Left and right structures were re-labeled with different identification numbers using the sagittal mid-line (in MNI space, x coordinate at zero) as a separator.
  m. **HO cort-only**. Same as the HO cortical-only atlas.
  n. **HO sym. cortical only**. HO atlas with only cortical regions. The symmetrical regions (the same region name on the contralateral hemisphere) are labeled with the *same* identification. Thus, this atlas is has *symmetrical* labels (e.g. both temporal pole regions are labeled with the same identification number). The default atlases given by FSL are symmetrical atlases.
  o. **HO subcortical-only**. HO atlas with only subcortical regions.
  p. **HO subcort-only**. Same as the HO subcortical-only atlas.
  q. **HO combined**. HO atlas with both cortical and sub-cortical regions. This atlas has non-symmetrical labeling (e.g. both temporal pole regions are labeled with a different identification number).
  r. **HO cortical + subcortical**. Same as the HO combined atlas.
  s. **JHU**. The Johns Hopkins University atlases. There are two white matter atlases: thee JHU labels and JHU tracts atlases.
  t. **MMP**. Multi-modal parcellation atlas. Sometimes referred to as the “Glasser Atlas” after the first author of the original publication.
  u. **Random atlas 10-10**,**000**. Atlases created with random parcels with a specified number of parcels (e.g. Random atlas 1,000 will have 1,000 parcels). These atlases were built in the ICBM 2009c Nonlinear Asymmetric template. Thus, these atlases are whole-brain atlases (includes cortical gray matter, subcortical gray matter, and white matter). See the ‘Atlases’ Methods section for more details.
  v. **Schaefer 100-1**,**000**. The Schaefer atlases with a specified number of parcels (e.g. Schaefer 100 will have 100 parcels). There are ten atlases of 100, 200, 300, 400, 500, 600, 700, 800, 900, and 1,000 parcels.
  w. **Yeo liberal**. The Yeo atlases where the boundaries of each parcel is extended slightly into the white matter, past the cortical boundary.
  x. **Yeo conservative**. The Yeo atlases where the boundaries of each parcel is extended slightly into the white matter, past the cortical boundary.
2. **Δ SFC**. The change in SFC between ictal and preictal stats (*SFC*_*ictal*_ *SFC*_*preictal*_). This indicates whether or not the change in functional connectivity is congruent with the underlying structural connectivity.
3. **Contact**. A single sensor on an electrode that records LFP. Not to be confused with an electrode. See Fig. S7, bottom.
4. **ECoG**: Electrocorticography.
5. **Electrode**. Not to be confused with contact. See Fig. S7, bottom.
6. **Derived atlas**: An atlas which was derived from another atlas. For example, the AAL 600 is derived from the AAL atlas in which its parcellations are further sub-divided using a specified algorithm. Derived atlases may also be sub-divided randomly so that it is both considered a random and derived atlas (a quasi-random atlas). The BNA is also a derived atlas in which it initially used the parcellations of the DK atlas.
7. **Functional connectivity (FC)**. The statistical relationship between two signals (two contacts in this study).
8. **grayordinate**. Atlas that includes gray matter structures, including cortical and subcortical gray matter regions.
9. **ROI**. Region of interest
10. **ROI, parcel, parcellation, region**. These terms may be used interchangeably in the literature. They refer to discrete areas of a brain. These regions are labeled with a categorical identification (rather than a continuous variable seen in templates - see Fig. S1), and all voxels or surface vertices with the same identification are part of thee same region.
11. **SEEG**: Stereoelectroeenccephalography.
12. **Structural connectivity (SC)**. The physical relationship between two brain regions. We use streamline counts in this manuscript from High Angular Resolution Diffusion Imaging.
13. **T1w**. T1-weighted MRI image.

---

**Fig. S1.**
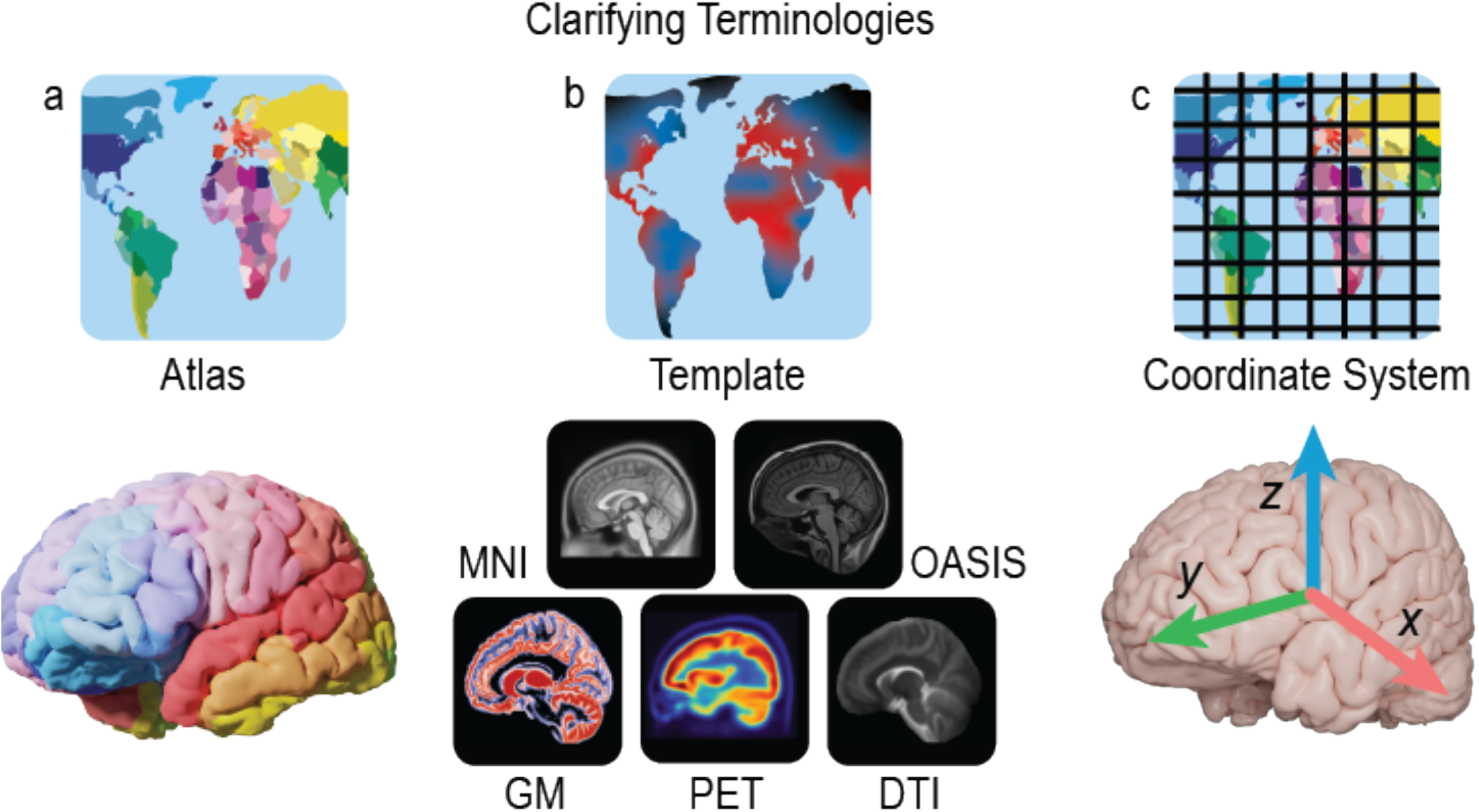
Atlas, Template, and Coordinate (Stereotactic) Space. These three terms are commonly confused in the neuroscience literature because they all relate to the “map” of the brain. “Atlas” and “template” are sometimes used interchangeably^3^, however, they are distinct. Here, we define them more formally. **a**, A brain ***atlas*** refers to a neurological map that defines brain region *labels*. We use this definition throughout the main text. **b**, An atlas is distinct from a brain ***template***, which refers to a brain *pattern*. Similar in common usage, a template is a mold, gauge, or starting point representation of the brain. Usually it is composed of multiple individuals’ brain representing an average of a population. Many templates exist and are reviewed in various publications^2,9^, The templates illustrated here are the MNI152 Nonlinear asymmetric 2009c T1w template (http://www.bic.mni.mcgill.ca), the OASIS brain template https://www.oasis-brains.org/ created and used by ANTs (http://stnava.github.io/ANTs/ with templates linked here), a gray matter probability map, a PET template, and a b0 DTI template. **c**, The coordinate system, or the **stereotactic space**, of the brain describes the physical positioning of the brain, similar to the geographical coordinate system of longitude and latitude of the Earth. Historically, a common stereotactic space was the Talairach space, and more recently, the MNI spaces. The analogy between the geographical terms of the Earth and the geographical terms of the brain is not exact. The analogy falls apart in that while there in one world, there are many brains. There is variability across populations and a spectrum of differences between species, therefore, it is challenging to represent one brain for use in every scientific study appropriately. **MNI**, Montreal Neurological Institute; **OASIS**, Open Access Series of Imaging Studies; **GM**, Gray Matter probability map; **PET**, Positron Emission Tomography; **DTI**, Diffusion Tensor Imaging.

**Table S1.**
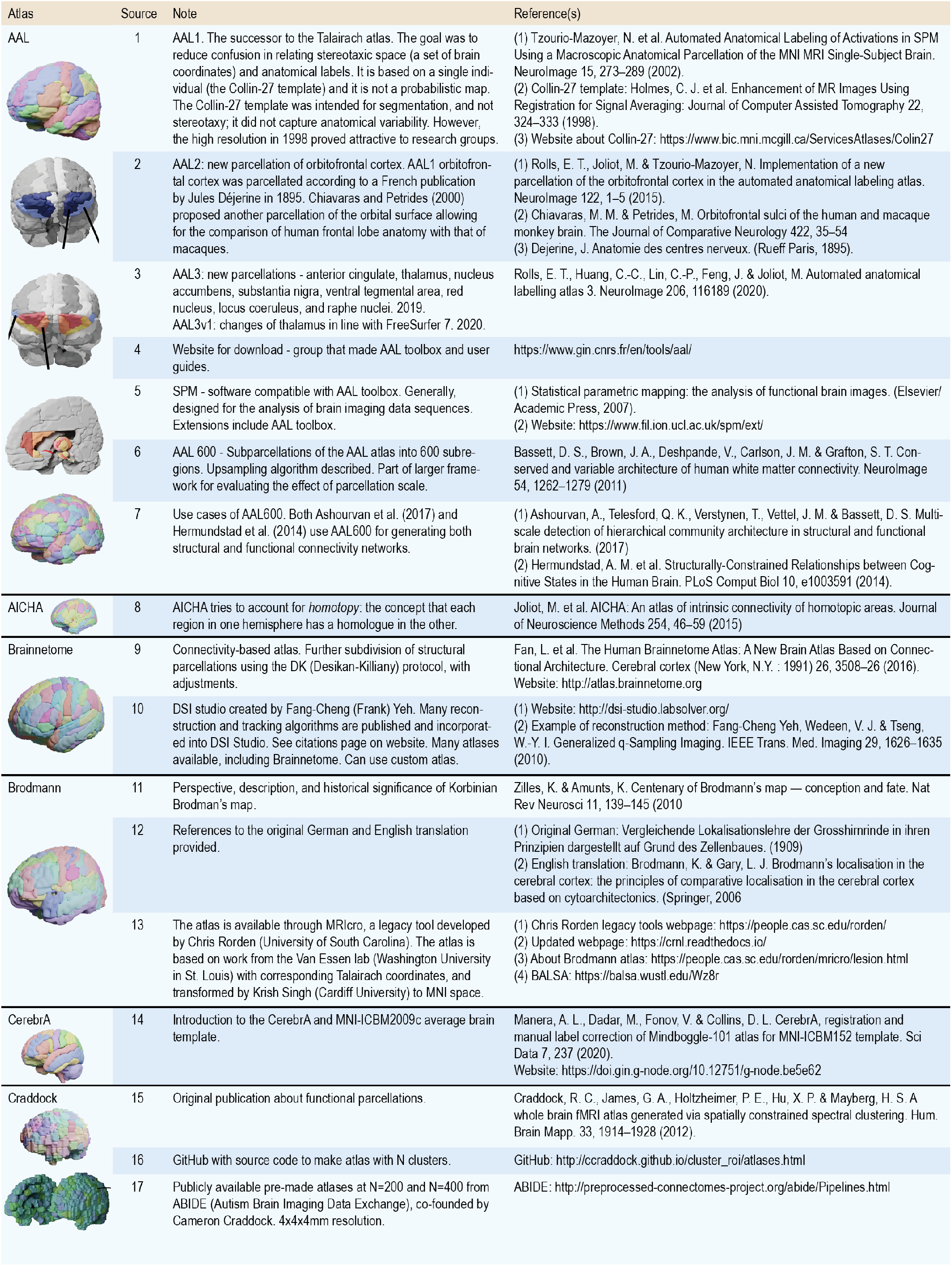
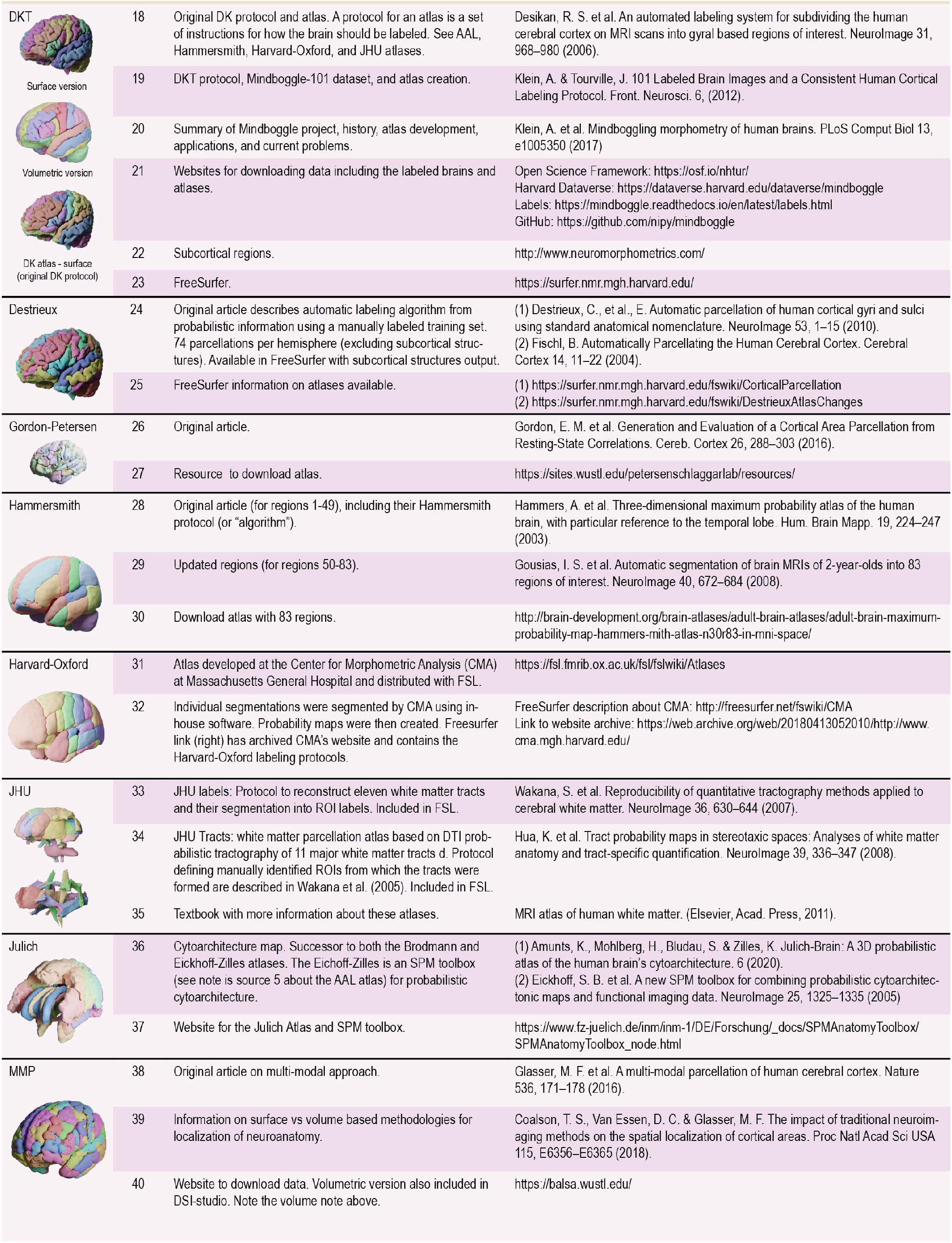
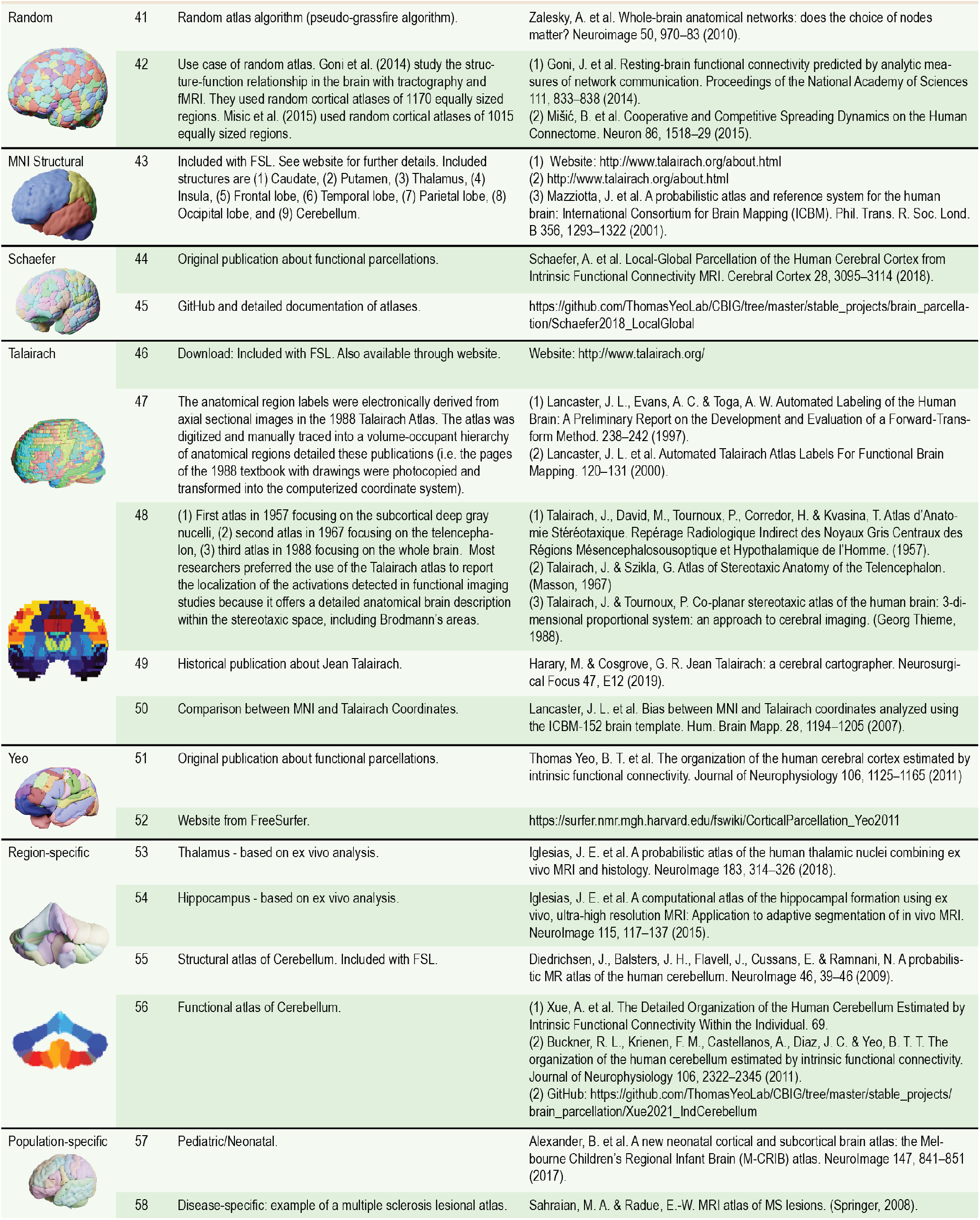
Atlas sources and references. This table provides a short note and references to the source material of common atlases in the neuroscience literature. See also Table 1.

**Fig. S2.**
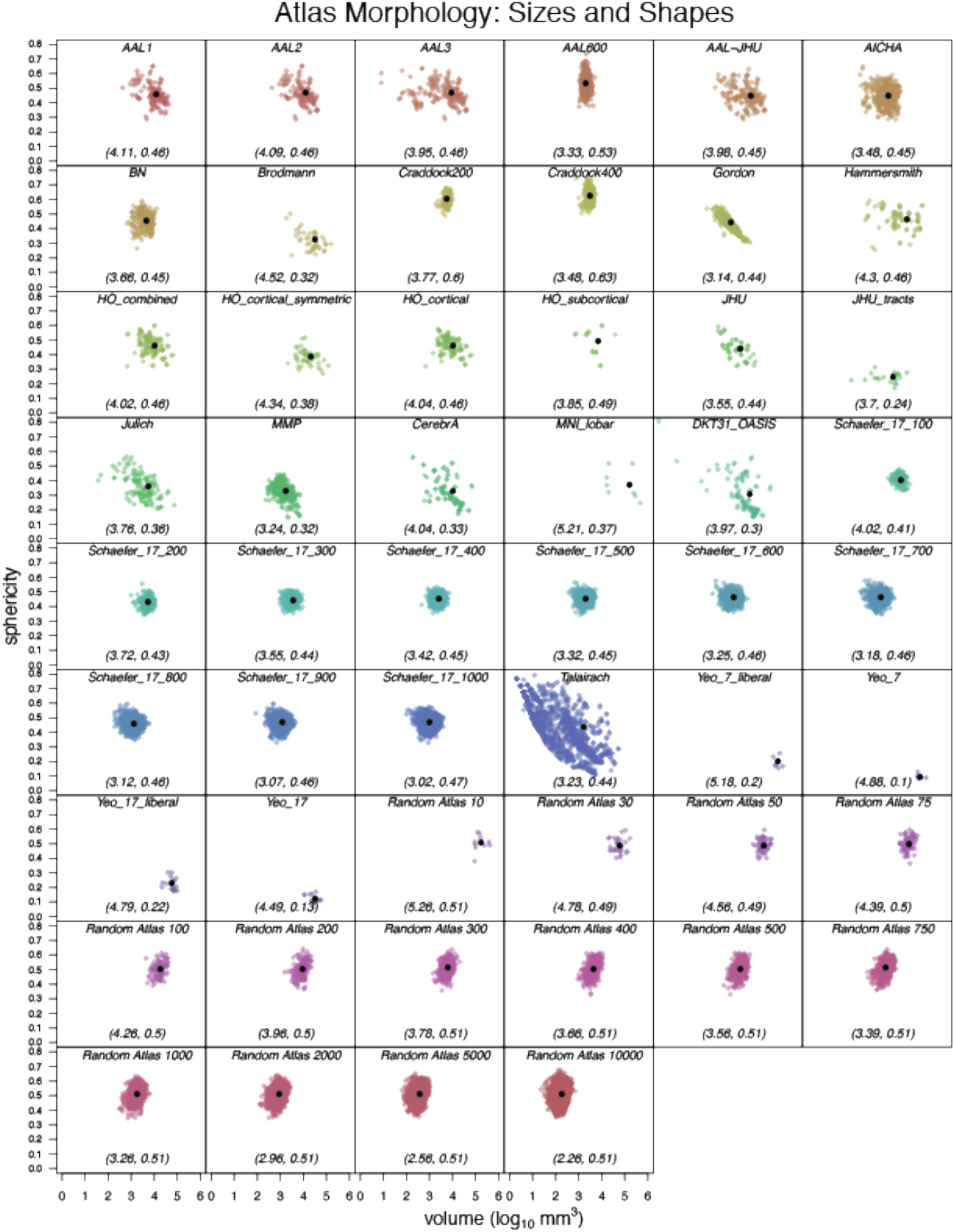
Atlas Morphology: Sizes and Shapes. All standard atlases and one permutation for each of the standard atlases are shown here. Volume means and sphericity means are in parentheses at the bottom of each graph. See Table S1 for atlas abbreviations, descriptions, and sources.

**Fig. S3.**
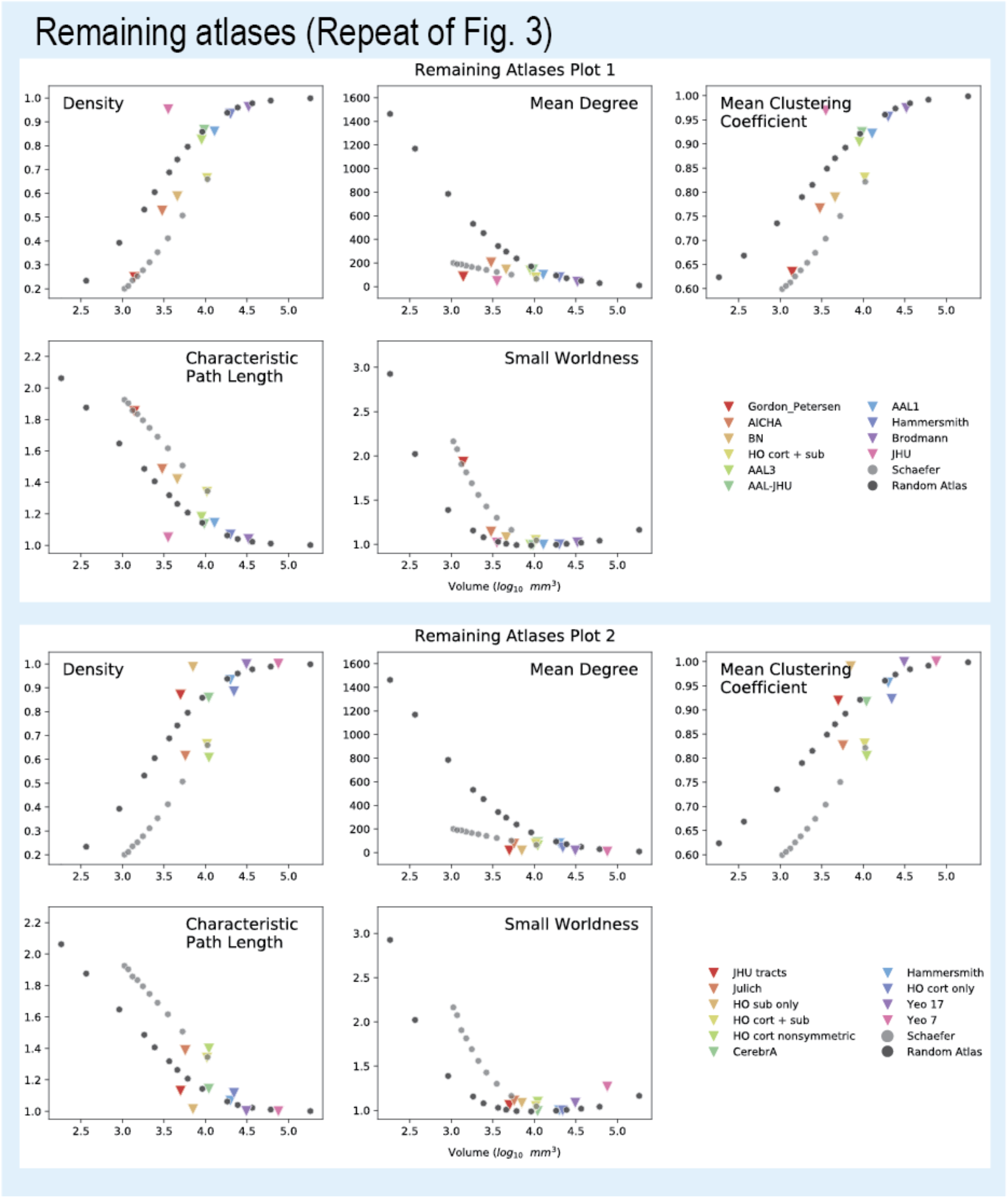
Structure-Function Correlation (SFC) for All Atlases. We show network measures the remaining atlases illustrated in Table 2. See Table S1 for atlas descriptions. **HO**, Harvard-Oxford; **Sub**, subcortical; **Cort**, cortical

**Fig. S4.**
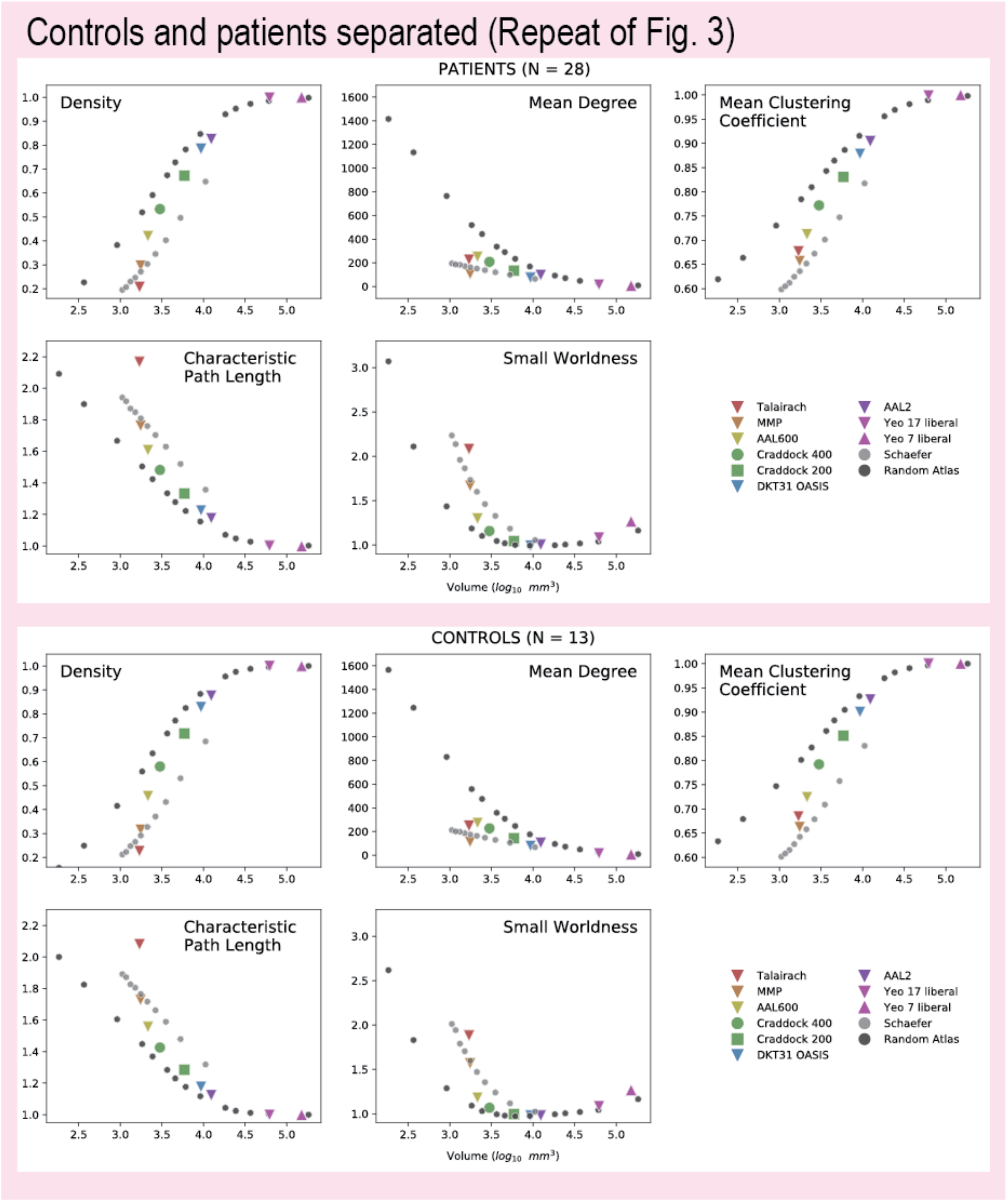
Network Measures: Controls vs Patients. We replicate Fig. 2 (N=41) in the manuscript by separating out controls (N=13) and patients (N=28). All global network measures above are similar between patients and controls, with patients having slightly lower (but not significant, Fig. 2 bottom right panel) measurements for the different network properties. Specific connectivity differences between controls and patients were not explored (e.g. to explore if connections from the hippocampus to the anterior cingulate are changed in temporal lobe epilepsy) and out of the scope of this manuscript. See Table S1 for atlas descriptions.

**Fig. S5.**
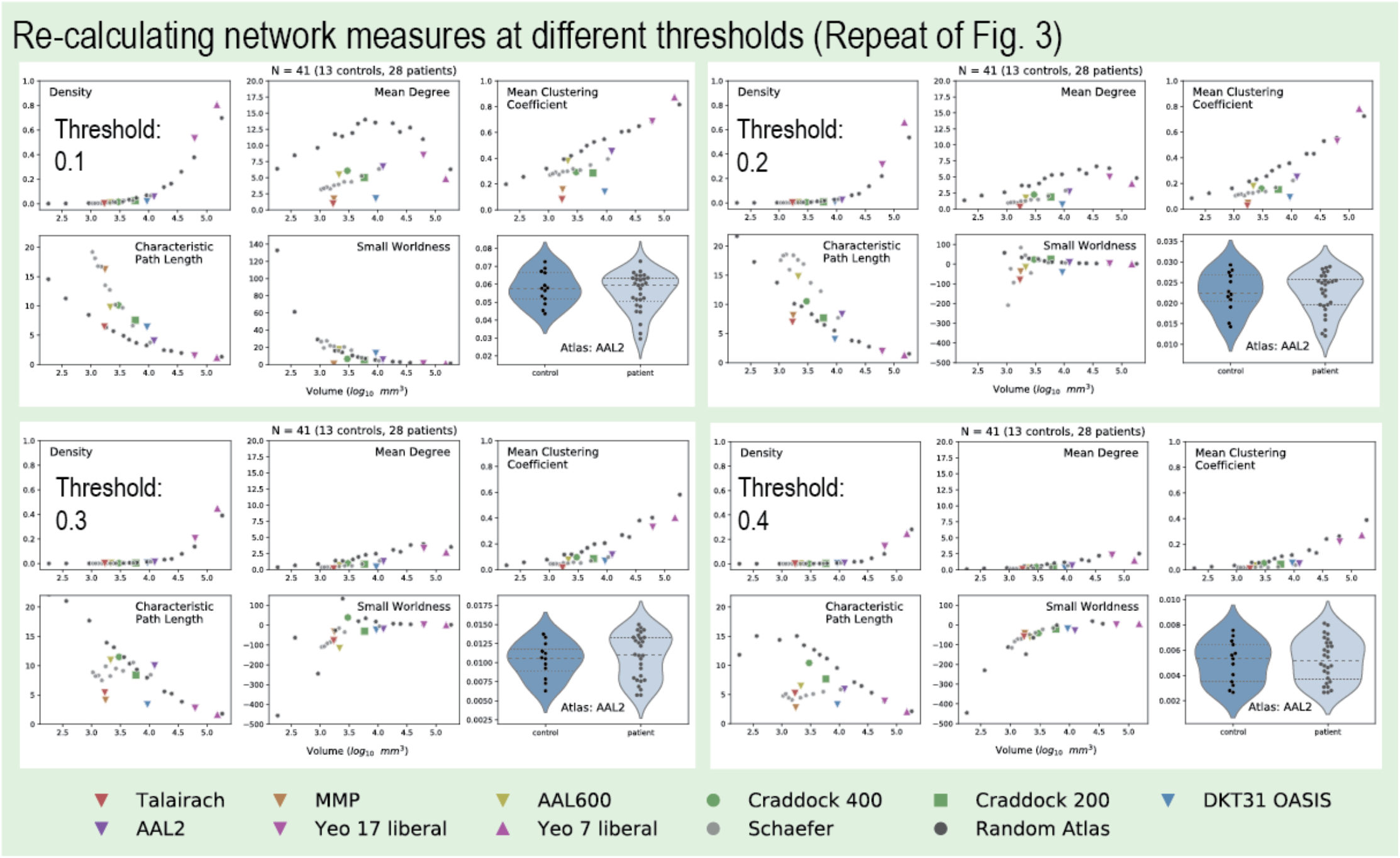
Network Measures: different thresholds. We replicate Fig. 2 (N=41) in the manuscript by calculating network measures using different thresholds. The main text figure includes all weights with no threshold (threshold = 0). We set thresholds at 01., 0.2, 0.3, and 0.4. This was done to show how various network measures may also change when eliminating low-level connections at different thresholds.

**Fig. S6.**
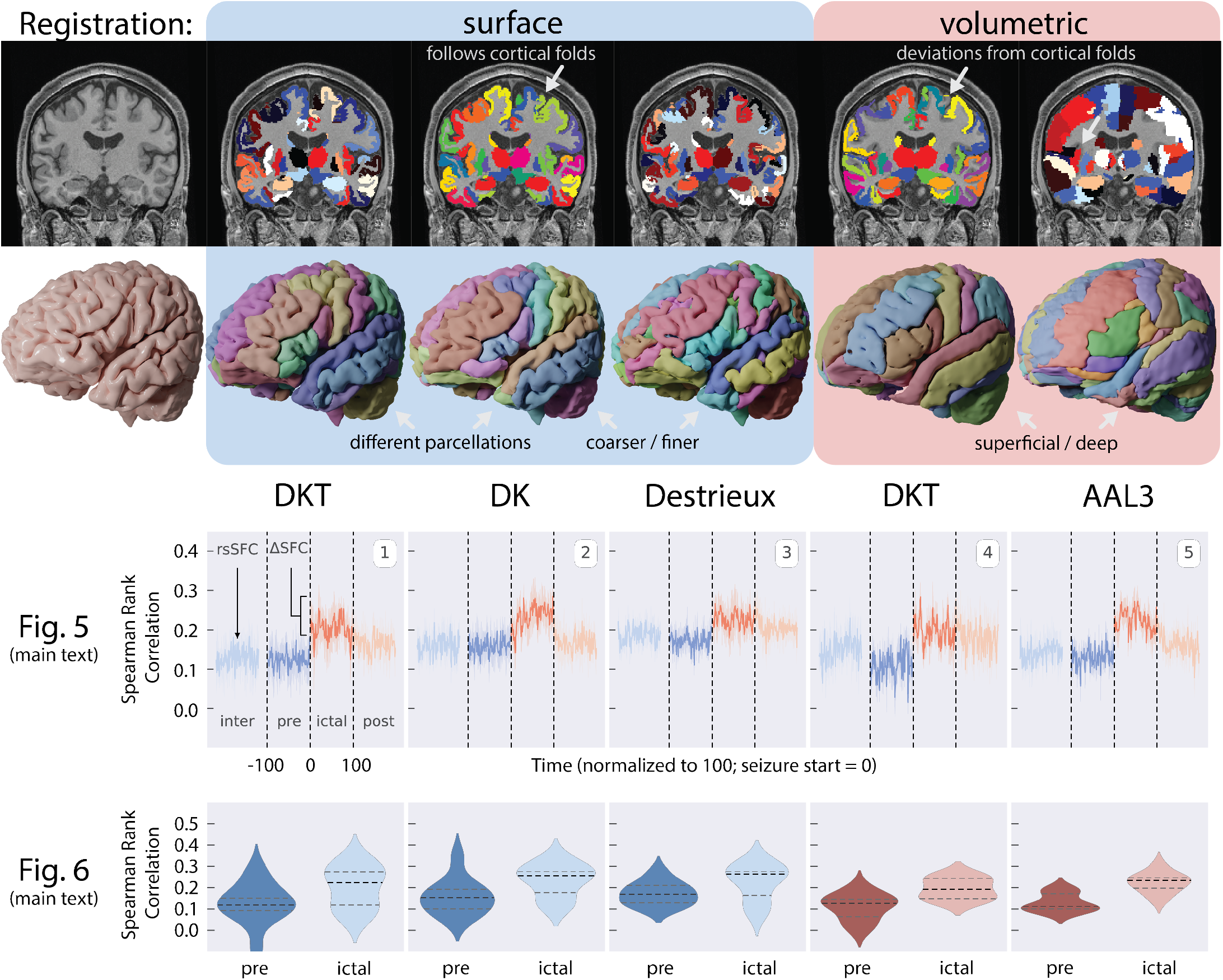
Effects of Registration: Volumetric- and Surface-based approaches. Volumetric-based analyses, as opposed to surface-based analyses, have been more prevalent in human neuroimaging studies for the last few decades^21^. Volumetric-based approaches to map the neocortex have been shown to be inaccurate in some cases. For example, the top row shows a single subject’s T1w image and the resulting labels of three atlases registered using a surface-based approach and two atlases using a volumetric-based approach. The DKT atlas using a surface-based approach follows the cortical folds of the T1w image closely, but the DKT atlas registered using a volumetric-based approach may have many mis-aligned areas. These images show the improved accuracy in mapping and labeling brain structures using surface-based analyses, but the adoption of surface-based analyses has been slow and attributed to five main reasons discussed in Coalson et. al 2018^21^. Briefly, it is due to (1) the need to compare results with existing volumetric-based studies, (2) the prevalence of volumetric-based tools compared to surface-based tools, (3) the learning curve of surface-based approaches; (4) an unawareness of the problems and benefits of each approach; (5) and uncertainty or skepticism as to how much of a difference these methodological choices make. In some cases, it may make a difference, however, it does not make a difference in this study. Here, we used a surface-based approach to register three different atlases to each patient. The atlases were outputs of FreeSurfer’s recon-all pipelinee^78^ - the DKT40, Desikan-Killiany (DK), and Destrieux atlases. The DKT atlas has a modified parcellations of the DK atlas, and the Destrieux atlas is an alternative atlas offered by the FreeSurfer piepline. The Destrieux atlas has a finer parcellation scheme (i.e., more number of regions). We repeat analyses of Fig. 5 and Fig. 6 of the main text, along with results from two volumetric-based atlases for side-by-side comparison. The volumetric-based atlases include the DKT (DKT31 OASIS) and AAL3 atlases. While the volumetric DKT atlas does not properly align and label the entire cortical gray matter regions, the AAL atlas extends deeply into the white matter and does label much of these gray matter regions. For the experimental design of this study in localizing electrode contacts and measuring structural connectivity, the AAL3 atlas provides the most power out of all these atlases in detecting a change in SFC. In the original AAL manuscript^88^, the authors “chose to extend the internal limit of the regions beyond the gray matter layer [to account for] anatomical variability”. This extension past the internal gray matter boundary may be optimal in our case for measuring SFC because the parcellations may capture streamlines that otherwise would have ended prematurely before reaching gray matter.

**Fig. S7.**
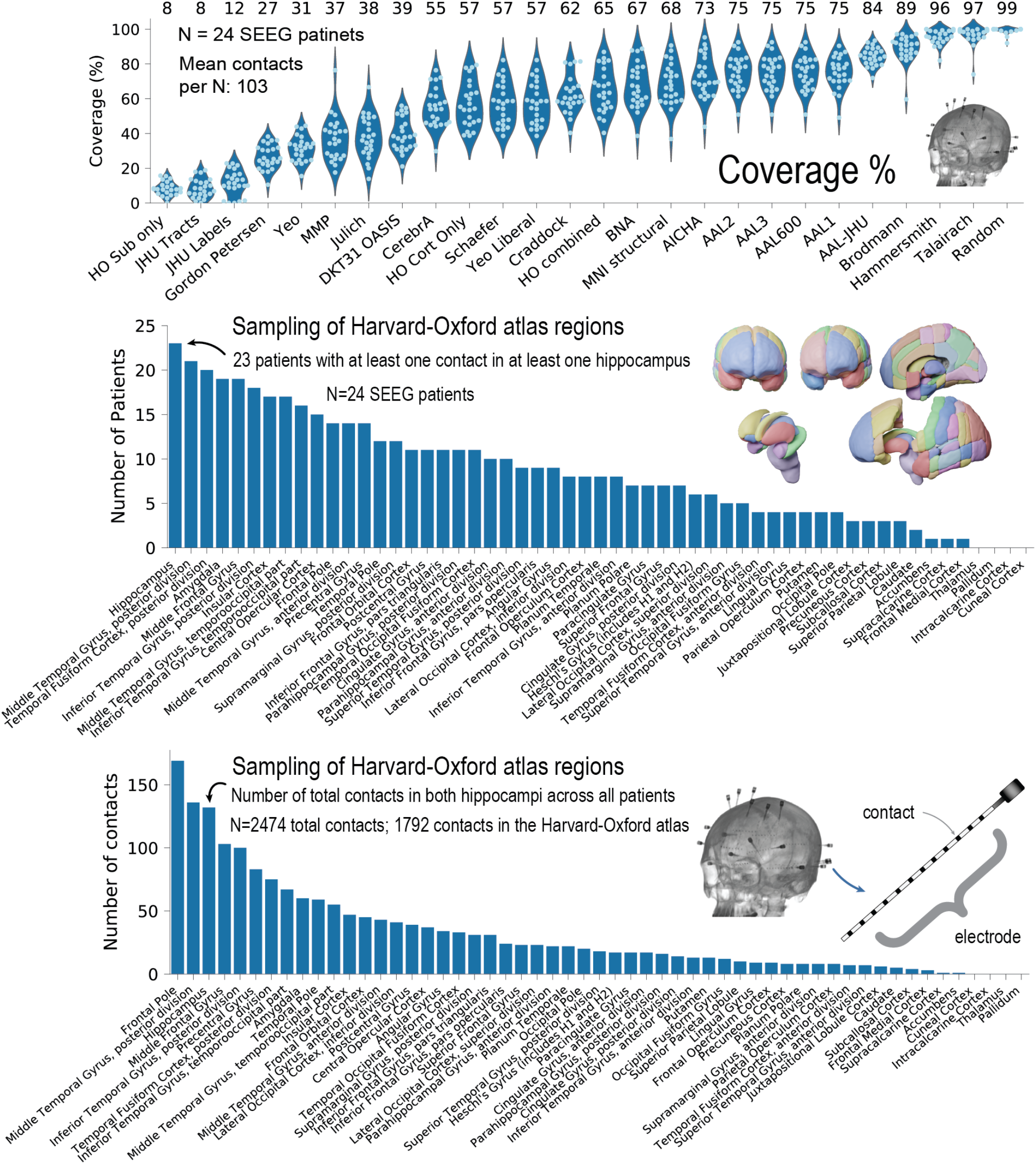
Coverage of electrode contacts. Top: We show the percentage of contacts assigned a region given an atlas. If a contact fell outside an atlas, it would not be assigned a location and would not be used in SFC analysis. We also show the Harvard-Oxford atlas regions (cortical and subcortical combined) that contain electrode contacts (middle and bottom figures). The middle figure shows the number of patients with at least one contact in an atlas region (at least one of the regions on both hemispheres). The bottom figure shows the total number of contacts in each listed region. Note that 1792 out of 2474 contacts (72%) contained within the brain parenchyma (gray matter or white matter) is higher than the mean percent coverage listed in the top figure (65% for the HO combined) because some patients with fewer contacts may have lower coverage by the atlas, thus bringing the mean percent down. Also note the larger number of contacts in the frontal pole because this region in the Harvard-Oxford atlas is large. We chose to show the Harvard-Oxford atlas because it has the largest effect size in Fig. 6.

**Fig. S8.**
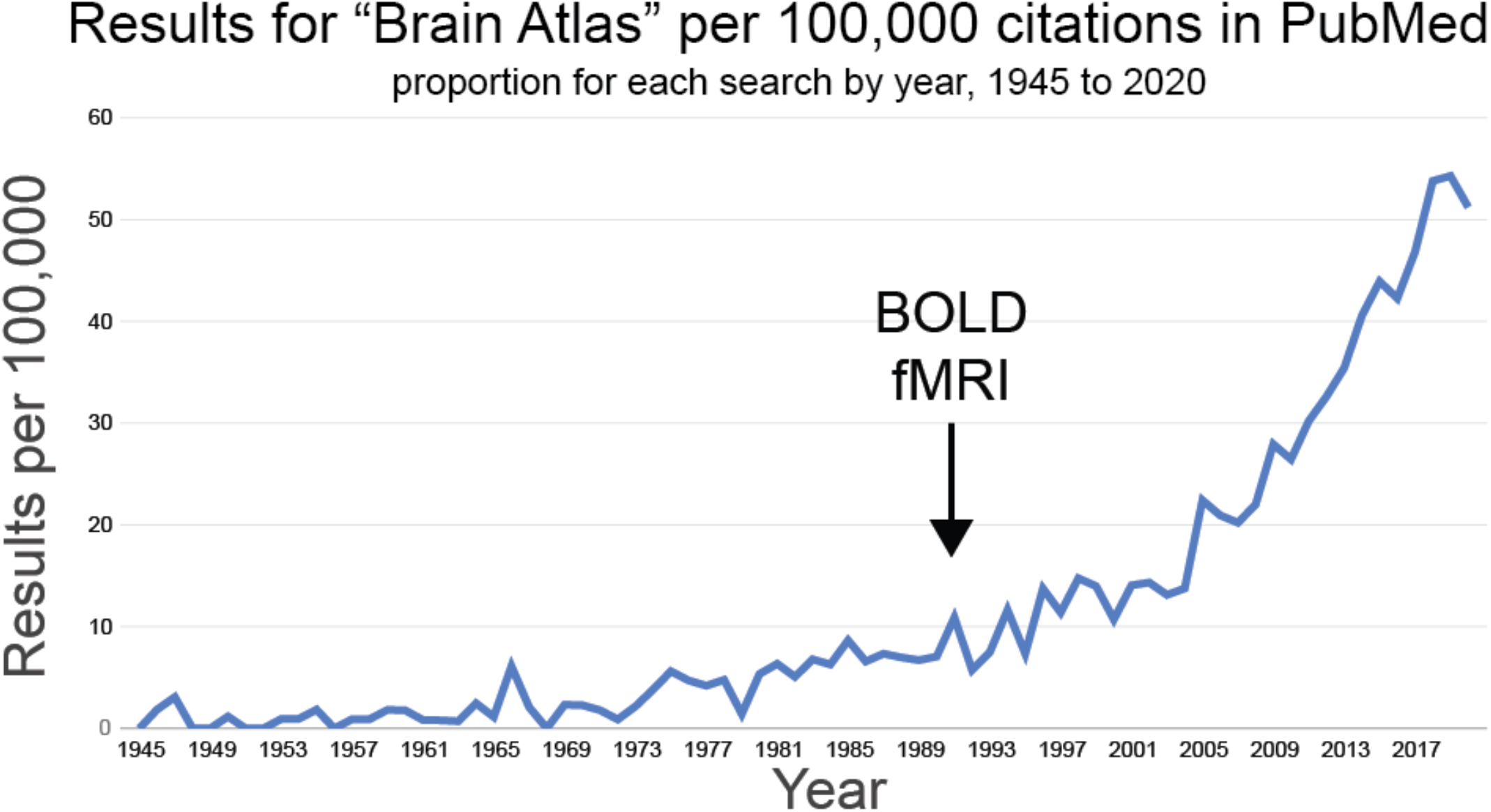
The increase in publications related to brain atlases. We searched for any publications since 1945 using the term “Brain Atlas” on PubMed. We note that since the introduction of BOLD fMRI in 1990, the need for neuroanatomical maps of the brain has increased, especially in the neuroimaging community. Many atlases have been published over the last 30 years, and many publications across the neuroscience literature have used these atlases. However, no comprehensive study exists evaluating, in any regard, to the suitability and nuances related to these atlases. We hope our work provides a valuable resource to others in our field, launches a larger discussion to critically evaluating the neuroanatomy of the brain, and direct future reproducible research for other scientists and clinician investigators.

**Fig. S9.**
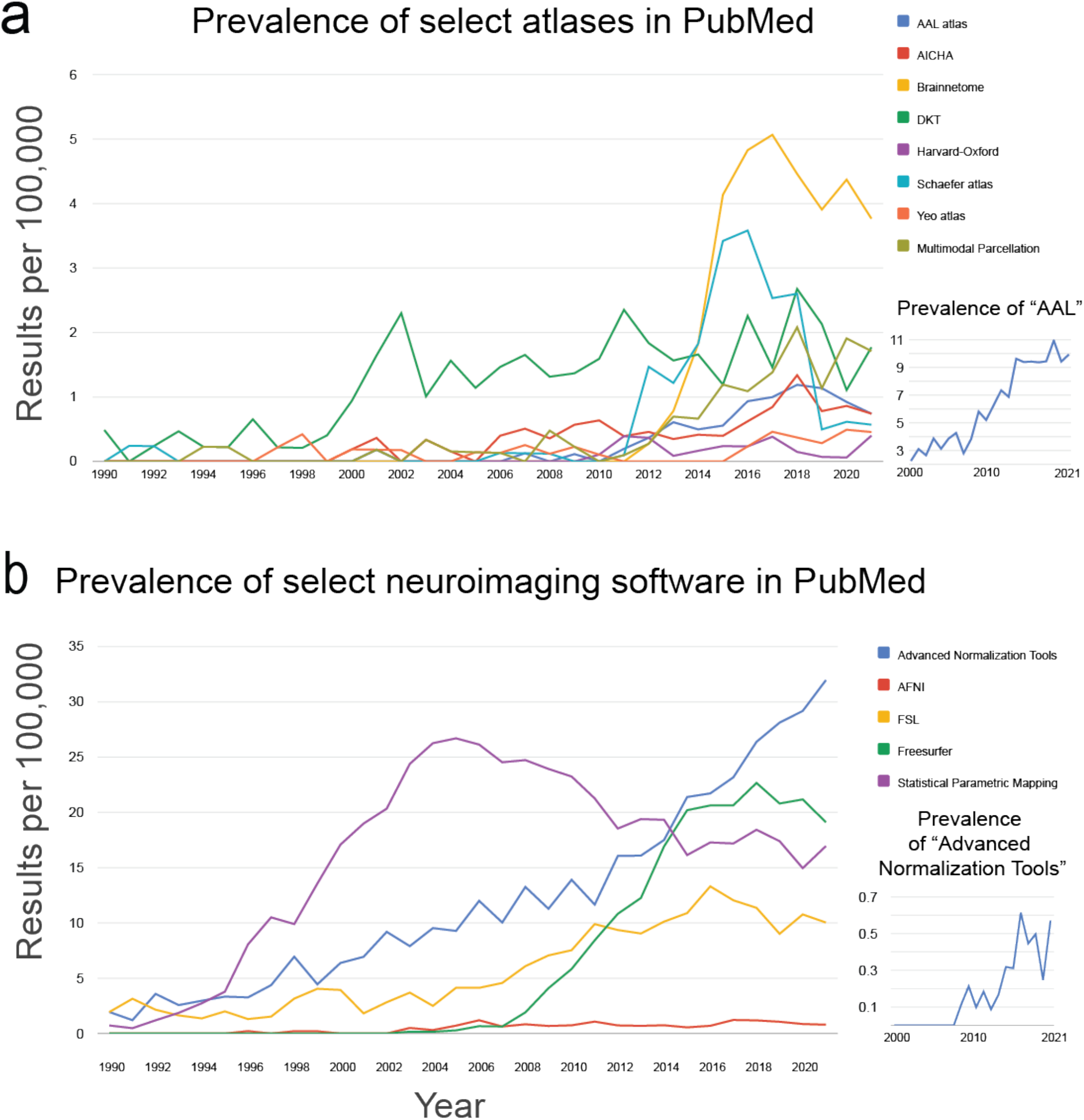
Prevalence of select brain atlases and neuroimaging software. **a**, We searched on PubMed for any publications since 1945 using the verbatim terms shown in each line graph legend. The tool used is from https://esperr.github.io/pubmed-by-year/^89^. This search was done to gain a better understanding how often the field is using different tools, and thus to make some recommendations as to which atlases to use and facilitating the comparison of results. Note that due to the prevalence of the term “AAL” which may not relate to the AAL atlas, we opted for the term “AAL atlas”. Another example is the use of “Multimodal Parcellation” rather than “MMP”. The search for “AAL” is shown at the bottom right, where articles appear before the original AAL manuscript in 2002^88^, most likely not relating to the AAL atlas. However, the prevalence of “AAL” increases substantially after 2002, more than other atlases. These search terms serves as a rough estimate of the prevalence of atlases, and may not reflect the true prevalence of each term. **b**, We show to prevalence of select neuroimaging software. Again, due to the ambiguity of search terms such as “ANTs”, we opted for the full name of the software, despite some manuscripts only having used the abbreviated terms. “Advanced normalization tools” searched in quotes is shown at the bottom right, having first appeared formally in the literature in 2009^90^.

**Fig. S10.**
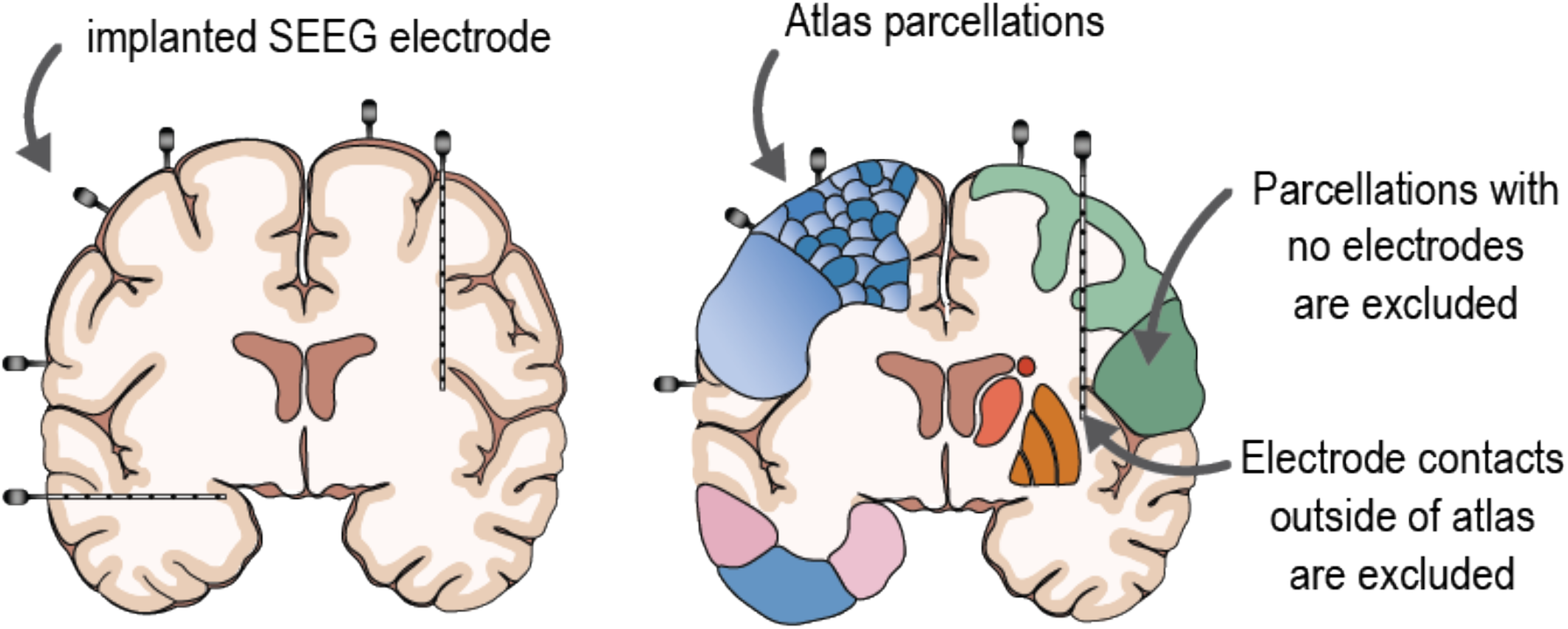
Electrode localization and region selection. Assignment of each electrode contact to an atlas regions was performed by rounding electrode coordinates (x,y,z) to the nearest voxel and indexing the given atlas at that voxel. Electrodes that fell outside the atlas of interest were excluded from subsequent analysis. The structural connectivity network, representing normalized streamline counts between each atlas region, was also down sampled to only include regions that contained at least one SEEG contact. This gave one static representation of structural connectivity. In the case where multiple electrodes fell in the same atlas ROI, a random electrode was selected to represent the functional activity of that neuroanatomically defined region.

**Table S2.** Patient and control demographics. Patient IDs with asterisk have clinically annotated seizures for structure-function calculation. Localization of the seizure onset zone was pulled from patient charts, either from the clinically hypothesized brain regions if the patient did not undergo surgery, or if the patient underwent surgery, the targeted location for resection or ablation. One control did not have age or sex information. **M**, Male; **F**: Female; **L**, left; **R**, Right; **NR**, Not reported

| Patient | Age | Sex | Localization: suspected seizure onset zone | Control | Age | Sex |
| --- | --- | --- | --- | --- | --- | --- |
| sub-patient01 | 58 | M | Poorly localized. R temporal interictal activity. | sub-control01 | 24 | M |
| sub-patient02 | 28 | F | L anterior temporal lobe | sub-control02 | 40 | F |
| sub-patient03 | 27 | F | L hippocampus and amygdala | sub-control03 | 31 | M |
| sub-patient04 | 20 | F | L basal ganglia infarct | sub-control04 | 29 | M |
| sub-patient05** | 36 | M | R frontal arteriovenous malformation | sub-control05 | 40 | M |
| sub-patient06 | 57 | F | Poorly localized. Possibly bitemporal onset | sub-control06 | 48 | F |
| sub-patient07** | 37 | M | L temporal lobe/hippocampus/amygdala | sub-control07 | 22 | M |
| sub-patient08** | 34 | M | R frontal, anterior cingulate gyrus | sub-control08 | 35 | F |
| sub-patient09** | 47 | F | L hippocampus | sub-control09 | 27 | F |
| sub-patient10 | 42 | F | R temporal lobe/L temporal lobe | sub-control10 | 67 | F |
| sub-patient11 | 27 | M | L hippocampus, then amygdala | sub-control11 | 33 | F |
| sub-patient12 | 35 | M | Poorly localized. Possibly multifocal epilepsy | sub-control12 | 27 | M |
| sub-patient13** | 36 | F | L temporal | sub-control13 | NR | NR |
| sub-patient14** | 29 | F | L superior Frontal Sulcus |  |  |  |
| sub-patient15 | 33 | F | L mesial temporal lobe |  |  |  |
| sub-patient16 | 29 | M | Poorly localized. Possibly multifocal epilepsy |  |  |  |
| sub-patient17 | 31 | F | L mesial temporal lobe |  |  |  |
| sub-patient18** | 26 | F | L heterotopia, left hippocampus |  |  |  |
| sub-patient19** | 23 | M | L temporal/posterior lateral neocortical |  |  |  |
| sub-patient20 | 30 | M | L temporal encephalomalacia |  |  |  |
| sub-patient21 | 24 | M | R anterior temporal lobe |  |  |  |
| sub-patient22 | 59 | F | R frontal-parietal lobe |  |  |  |
| sub-patient23 | 28 | F | L or R superior temporal gyrus |  |  |  |
| sub-patient24** | 47 | F | R anterior temporal |  |  |  |
| sub-patient25 | 40 | F | L temporal lobe near Heschl's gyrus |  |  |  |
| sub-patient26 | 37 | F | L amygdala/anterior temporal pole |  |  |  |
| sub-patient27 | 30 | M | L amygdala/hippocampus |  |  |  |
| sub-patient28** | 28 | M | L mesial temporal lobe |  |  |  |

## References

[1] Klein, A. & Tourville, J. 101 Labeled Brain Images and a Consistent Human Cortical Labeling Protocol. Frontiers in Neuroscience 6 (2012).

[2] Mandal, P. K., Mahajan, R. & Dinov, I. D. Structural brain atlases: design, rationale, and applications in normal and pathological cohorts. Journal of Alzheimer’s disease : JAD 31 Suppl 3, S169–88 (2012).

[3] Dickie, D. A. et al. Whole Brain Magnetic Resonance Image Atlases: A Systematic Review of Existing Atlases and Caveats for Use in Population Imaging. Front Neuroinform 11, 1 (2017).

[4] Bohland, J. W., Bokil, H., Allen, C. B. & Mitra, P. P. The Brain Atlas Concordance Problem: Quantitative Comparison of Anatomical Parcellations. PLoS ONE 4, e7200 (2009).

[5] Beal, D. S. et al. The trajectory of gray matter development in Broca’s area is abnormal in people who stutter. Frontiers in Human Neuroscience 9 (2015).

[6] Van Horn, J. D. et al. Mapping Connectivity Damage in the Case of Phineas Gage. PLoS ONE 7, e37454 (2012).

[7] Barker, F. G. Phineas among the phrenologists: the American crowbar case and nineteenthcentury theories of cerebral localization. Journal of Neurosurgery 82, 672–682 (1995).

[8] Mazziotta, J. et al. A probabilistic atlas and reference system for the human brain: International Consortium for Brain Mapping (ICBM). Philosophical Transactions of the Royal Society of London. Series B: Biological Sciences 356, 1293–1322 (2001).

[9] Evans, A. C., Janke, A. L., Collins, D. L. & Baillet, S. Brain templates and atlases. NeuroImage 62, 911–22 (2012).

[10] National Geographic Society Encyclopedic entry. Atlas (2022). URL https://www.nationalgeographic.org/encyclopedia/atlas/.

[11] National Geographic Society Encyclopedic entry. Border (2022). URL https://www.nationalgeographic.org/encyclopedia/border/.

[12] Van Essen, D. C. et al. The WU-Minn Human Connectome Project: an overview. NeuroImage 80, 62–79 (2013).

[13] Shah, P. et al. Characterizing the role of the structural connectome in seizure dynamics. Brain: A Journal of Neurology (2019).

[14] Ashourvan, A. et al. Pairwise maximum entropy model explains the role of white matter structure in shaping emergent co-activation states. Commun Biol 4 (2021).

[15] Shmueli, G. To Explain or to Predict. Statistical Science 25 (2010).

[16] Makris, N. et al. Decreased volume of left and total anterior insular lobule in schizophrenia. Schizophrenia Research 83, 155–171 (2006).

[17] Hammers, A. et al. Three-dimensional maximum probability atlas of the human brain, with particular reference to the temporal lobe. Human Brain Mapping 19, 224–247 (2003).

[18] Thomas Yeo, B. T. et al. The organization of the human cerebral cortex estimated by intrinsic functional connectivity. J Neurophysiol 106, 1125–65 (2011).

[19] Schaefer, A. et al. Local-Global Parcellation of the Human Cerebral Cortex from Intrinsic Functional Connectivity MRI. Cerebral Cortex 28, 3095–3114 (2018).

[20] Glasser, M. F. et al. A multi-modal parcellation of human cerebral cortex. Nature 536, 171–178 (2016).

[21] Coalson, T. S., Van Essen, D. C. & Glasser, M. F. The impact of traditional neuroimaging methods on the spatial localization of cortical areas. Proceedings of the National Academy of Sciences 115, E6356–E6365 (2018).

[22] Wu, Z., Xu, D., Potter, T., Zhang, Y. & The, A. D. N. I. Effects of Brain Parcellation on the Characterization of Topological Deterioration in Alzheimer’s Disease. Frontiers in Aging Neuroscience 11, 113 (2019).

[23] Proix, T., Bartolomei, F., Guye, M. & Jirsa, V. K. Individual brain structure and modelling predict seizure propagation. Brain : a journal of neurology 140, 641–654 (2017).

[24] Wirsich, J. et al. Whole-brain analytic measures of network communication reveal increased structure-function correlation in right temporal lobe epilepsy. NeuroImage. Clinical 11, 707–718 (2016).

[25] Willner, P. The validity of animal models of depression. Psychopharmacology 83, 1–16 (1984).

[26] Bassett, D. S., Zurn, P. & Gold, J. I. On the nature and use of models in network neuroscience. Nature Reviews Neuroscience 19, 566–578 (2018).

[27] Association, A. P. Technical recommendations for psychological tests and diagnostic techniques. Psychological bulletin 51, 1–38 (1954).

[28] Salehi, M. et al. There is no single functional atlas even for a single individual: Functional parcel definitions change with task. NeuroImage 208, 116366 (2020).

[29] Albers, K. J. et al. Using connectomics for predictive assessment of brain parcellations. NeuroImage 238, 118170 (2021).

[30] Zalesky, A. et al. Whole-brain anatomical networks: does the choice of nodes matter? Neuroimage 50, 970–83 (2010).

[31] Cocchi, L. et al. Disruption of structure-function coupling in the schizophrenia connectome. NeuroImage. Clinical 4, 779–87 (2014).

[32] Sathian, K. & Crosson, B. Structure-function correlations in stroke. Neuron 85, 887–9 (2015).

[33] Gorgolewski, K. J. et al. http://NeuroVault.org: a web-based repository for collecting and sharing unthresholded statistical maps of the human brain. Frontiers in neuroinformatics 9, p8 (2015).

[34] Lawrence, R. M. et al. Standardizing human brain parcellations. Scientific data 8, 78 (2021).

[35] Alexander, B. et al. A new neonatal cortical and subcortical brain atlas: the Melbourne Children’s Regional Infant Brain (M-CRIB) atlas. NeuroImage 147, 841–851 (2017).

[36] Brennan, B. P. et al. Use of an Individual-Level Approach to Identify Cortical Connectivity Biomarkers in Obsessive-Compulsive Disorder. Biological psychiatry. Cognitive neuroscience and neuroimaging 4, 27–38 (2019).

[37] Cabezas, M., Oliver, A., Lladó, X., Freixenet, J. & Cuadra, M. B. A review of atlas-based segmentation for magnetic resonance brain images. Computer methods and programs in biomedicine 104, e158–77 (2011).

[38] Caspers, S., Eickhoff, S. B., Zilles, K. & Amunts, K. Microstructural grey matter parcellation and its relevance for connectome analyses. NeuroImage 80, 18–26 (2013).

[39] Diedrichsen, J., Balsters, J. H., Flavell, J., Cussans, E. & Ramnani, N. A probabilistic MR atlas of the human cerebellum. NeuroImage 46, 39–46 (2009).

[40] Belzung, C. & Lemoine, M. Criteria of validity for animal models of psychiatric disorders: focus on anxiety disorders and depression. Biology of mood & anxiety disorders 1, 9 (2011).

[41] Association, A. E. R. (ed.) Standards for Educational and Psychological Testing (American Educational Research Association, Lanham, MD, 2014).

[42] Sporns, O., Tononi, G. & Kötter, R. The human connectome: A structural description of the human brain. PLoS computational biology 1, e42 (2005).

[43] Fornito, A., Zalesky, A. & Bullmore, E. Fundamentals of Brain Network Analysis (Academic Press, 2016).

[44] Sporns, O. The human connectome: a complex network. Annals of the New York Academy of Sciences 1224, 109–125 (2011).

[45] Greene, P., Li, A., González-Martínez, J. & Sarma, S. V. Classification of Stereo-EEG Contacts in White Matter vs. Gray Matter Using Recorded Activity. Frontiers in Neurology 11 (2021).

[46] Mercier, M. R. et al. Evaluation of cortical local field potential diffusion in stereotactic electroencephalography recordings: A glimpse on white matter signal. NeuroImage 147, 219–232 (2017).

[47] Young, J. J. et al. Quantitative Signal Characteristics of Electrocorticography and Stereoelectroencephalography: The Effect of Contact Depth. Journal of clinical neurophysiology : official publication of the American Electroencephalographic Society 36, 195–203 (2019).

[48] Revell, A. Y. et al. White Matter Signals Reflect Information Transmission Between Brain Regions During Seizures. bioRxiv (2021).

[49] Henderson, M. X. et al. Spread of a-synuclein pathology through the brain connectome is modulated by selective vulnerability and predicted by network analysis. Nat Neurosci 22, 1248–1257 (2019).

[50] Wang, H. E. et al. VEP atlas: An anatomic and functional human brain atlas dedicated to epilepsy patients. J Neurosci Methods 348, 108983 (2021).

[51] Doucet, G. E. et al. Atlas55+: Brain Functional Atlas of Resting-State Networks for Late Adulthood. Cereb Cortex 31, 1719–1731 (2021).

[52] Muñoz-Castañeda, R. et al. Cellular anatomy of the mouse primary motor cortex. Nature 598, 159–166 (2021).

[53] Huang, C.-C., Rolls, E. T., Feng, J. & Lin, C.-P. An extended Human Connectome Project multimodal parcellation atlas of the human cortex and subcortical areas. Brain Structure and Function (2021).

[54] Syversen, I. F. et al. Structural connectivity-based segmentation of the human entorhinal cortex. NeuroImage 245, 118723 (2021).

[55] Zhu, J. et al. Integrated structural and functional atlases of Asian children from infancy to childhood. NeuroImage 245, 118716 (2021).

[56] Joglekar, A. et al. A spatially resolved brain region-and cell type-specific isoform atlas of the postnatal mouse brain. Nat Commun 12, 463 (2021).

[57] Callaway, E. M. et al. A multimodal cell census and atlas of the mammalian primary motor cortex. Nature 598, 86–102 (2021).

[58] Lewis, J. D., Bezgin, G., Fonov, V. S., Collins, D. L. & Evans, A. C. A sub+cortical fMRI-based surface parcellation. Human Brain Mapping (2021).

[59] Gordon, E. M. et al. Generation and Evaluation of a Cortical Area Parcellation from Resting-State Correlations. Cerebral Cortex Cereb. Cortex 26, 288–303 (2016).

[60] Sinha, N. et al. Focal to bilateral tonic–clonic seizures are associated with widespread network abnormality in temporal lobe epilepsy. Epilepsia 62, 729–741 (2021). URL https://onlinelibrary.wiley.com/doi/10.1111/epi.16819.

[61] Royer, J. et al. An Open MRI Dataset for Multiscale Neuroscience. preprint, Neuroscience (2021). URL http://biorxiv.org/lookup/doi/10.1101/2021.08.04.454795.

[62] Fan, L. et al. The Human Brainnetome Atlas: A New Brain Atlas Based on Connectional Architecture. Cerebral cortex (New York, N.Y. : 1991) 26, 3508–26 (2016).

[63] Perlaki, G. et al. Comparison of accuracy between FSL’s FIRST and Freesurfer for caudate nucleus and putamen segmentation. Scientific Reports 7, 2418 (2017). URL http://www.nature.com/articles/s41598-017-02584-5.

[64] Mišić, B. et al. Cooperative and Competitive Spreading Dynamics on the Human Connectome. Neuron 86, 1518–29 (2015).

[65] Betzel, R. F. et al. Structural, geometric and genetic factors predict interregional brain connectivity patterns probed by electrocorticography. Nature Biomedical Engineering (2019).

[66] Smith, R. E., Tournier, J. D., Calamante, F. & Connelly, A. SIFT2: Enabling dense quantitative assessment of brain white matter connectivity using streamlines tractography. Neuroimage 119, 338–51 (2015).

[67] Bijsterbosch, J., Smith, S. M. & Beckmann, C. F. An Introduction to Resting State FMRI Functional Connectivity (Oxford University Press, 2017).

[68] Wagenaar, J. B., Brinkmann, B. H., Ives, Z., Worrell, G. A. & Litt, B. A multimodal platform for cloud-based collaborative research (IEEE, 2013).

[69] Kini, L. G., Davis, K. A. & Wagenaar, J. B. Data integration: Combined imaging and electro-physiology data in the cloud. NeuroImage 124, 1175–1181 (2016).

[70] Cieslak, M. et al. QSIPrep: an integrative platform for preprocessing and reconstructing diffusion MRI data. Nat Methods 18, 775–778 (2021).

[71] Fang-Cheng, Y., Wedeen, V. J. & Tseng, W.-Y. I. Generalized q-Sampling Imaging. IEEE Transactions on Medical Imaging 29, 1626–1635 (2010).

[72] Otsu, N. A Threshold Selection Method from Gray-Level Histograms. IEEE Transactions on Systems, Man, and Cybernetics 9, 62–66 (1979).

[73] Bonilha, L., Gleichgerrcht, E., Nesland, T., Rorden, C. & Fridriksson, J. Gray Matter Axonal Connectivity Maps. Frontiers in Psychiatry 6 (2015).

[74] Park, B., Eo, J. & Park, H.-J. Structural Brain Connectivity Constrains within-a-Day Variability of Direct Functional Connectivity. Frontiers in Human Neuroscience 11, 408 (2017).

[75] Taylor, P. N. et al. The impact of epilepsy surgery on the structural connectome and its relation to outcome. NeuroImage. Clinical 18, 202–214 (2018).

[76] Fonov, V. et al. Unbiased average age-appropriate atlases for pediatric studies. NeuroImage 54, 313–327 (2011).

[77] Jenkinson, M., Beckmann, C. F., Behrens, T. E. J., Woolrich, M. W. & Smith, S. M. FSL. NeuroImage 62, 782–790 (2012).

[78] Dale, A. M., Fischl, B. & Sereno, M. I. Cortical Surface-Based Analysis. NeuroImage 9, 179–194 (1999). URL https://linkinghub.elsevier.com/retrieve/pii/S1053811998903950.

[79] Maslov, S. Specificity and Stability in Topology of Protein Networks. Science 296, 910–913 (2002).

[80] Litt, B. et al. Epileptic Seizures May Begin Hours in Advance of Clinical Onset. Neuron 30, 51–64 (2001).

[81] Azarion, A. A. et al. An open-source automated platform for three-dimensional visualization of subdural electrodes using CT-MRI coregistration. Epilepsia 55, 2028–2037 (2014).

[82] Ludwig, K. A. et al. Using a common average reference to improve cortical neuron recordings from microelectrode arrays. Journal of neurophysiology 101, 1679–89 (2009).

[83] Kramer, M. A. et al. Coalescence and fragmentation of cortical networks during focal seizures. The Journal of neuroscience : the official journal of the Society for Neuroscience 30, 10076–85 (2010).

[84] Khambhati, A. N. et al. Dynamic Network Drivers of Seizure Generation, Propagation and Termination in Human Neocortical Epilepsy. PLoS computational biology 11, e1004608 (2015).

[85] Khambhati, A. N., Davis, K. A., Lucas, T. H., Litt, B. & Bassett, D. S. Virtual Cortical Resection Reveals Push-Pull Network Control Preceding Seizure Evolution. Neuron 91, 1170–1182 (2016).

[86] Khambhati, A. N. et al. Recurring Functional Interactions Predict Network Architecture of Interictal and Ictal States in Neocortical Epilepsy. eNeuro 4, ENEURO.0091–16.2017 (2017).

[87] Newson, J. J. & Thiagarajan, T. C. EEG Frequency Bands in Psychiatric Disorders: A Review of Resting State Studies. Frontiers in Human Neuroscience 12, 521 (2019).

[88] Tzourio-Mazoyer, N. et al. Automated anatomical labeling of activations in SPM using a macroscopic anatomical parcellation of the MNI MRI single-subject brain. NeuroImage 15, 273–89 (2002).

[89] Sperr, E. PubMed by Year (2022). URL https://esperr.github.io/pubmed-by-year/.

[90] Avants, B. B. et al. The optimal template effect in hippocampus studies of diseased populations. NeuroImage 49, 2457–2466 (2010). URL https://linkinghub.elsevier.com/retrieve/pii/S1053811909010611.

